# IL-21-producing peripheral helper T cells associate with autoimmune bile duct injury in biliary atresia

**DOI:** 10.64898/2026.07.08.736942

**Authors:** Mengting Liu, Weida Meng, Yuke Chen, Shaojing Wu, Manning Qian, Di Chen, Jie Zhang, Jianxiang Dong, Yifan Yang, Jingying Jiang, Teng Li, Qi Shi, Xufang Gu, Shaoyang Sun, Wenqing Qiu, Rui Dong, Xiaoming Zhang, Shan Zheng, Gong Chen, Yun Liu

**Author notes:** **Correspondence to**: Yun Liu, PhD,; Gong Chen, MD & PhD,; or Shan Zheng, MD & PhD. These authors contributed equally to this work (Mengting Liu, Weida Meng).

## Abstract

**Background:** Biliary atresia (BA) is a severe neonatal liver disease characterized by progressive fibrosis and bile duct obliteration.

**Objective:** Although immune dysregulation is implicated in the pathogenesis of BA, the specific mechanisms driving bile duct injury remain incompletely understood. This study aimed to characterize tertiary lymphoid structures (TLSs) within extrahepatic biliary remnants (EBRs), identify their cellular mediators, and evaluate the therapeutic potential of targeting IL-21 receptor signaling.

**Design:** We performed integrated bulk RNA sequencing, single-cell RNA sequencing, spatial transcriptomics, multiplex immunohistochemistry, and flow cytometry on clinical samples from BA patients and non-BA cholestatic controls. TLS maturation was assessed by CD23 immunohistochemistry in EBRs from 148 BA patients and correlated with clinical parameters. Anti-IL-21R antibody treatment was evaluated in a rhesus rotavirus-induced BA mouse model, with treatment initiated on day 4 post-infection.

**Results:** TLSs were identified in BA EBRs with significantly higher prevalence than in matched liver tissues. Mature TLSs containing CD23⁺ germinal centers were associated with elevated serum matrix metalloproteinase-7, more advanced hepatic fibrosis, and localized autoantibody deposition on injured bile ducts. Single-cell profiling revealed expanded CD4^+^ T peripheral helper (Tph) cells expressing IL-21 and CXCL13 within TLS-containing EBRs. Tph cells were enriched in peripheral blood of BA patients compared to non-BA cholestatic controls (*P* = 0.0025), and serum IL-21 was significantly elevated (*P* < 0.0001). Post-infection IL-21R blockade in the mouse model reduced jaundice incidence, improved weight gain, prevented extrahepatic biliary obstruction, and significantly improved long-term survival.

**Conclusion:** TLSs in BA extrahepatic biliary remnants harbor expanded Tph cells associated with IL-21-mediated B cell activation and bile duct injury. IL-21R blockade ameliorated disease in a murine BA model, identifying the IL-21/IL-21R axis as a potential therapeutic target warranting further investigation.

**Key Messages:** *What is already known on this topic:* Immune dysregulation contributes to biliary atresia (BA), with documented lymphocyte infiltration and defective B cell tolerance. However, the cellular mechanisms linking local immune activation to bile duct injury are unclear, and the roles of organized lymphoid structures and specific CD4⁺ T cell subsets in orchestrating local humoral responses have not been characterized.

*What this study adds:* This study demonstrates that mature tertiary lymphoid structures in extrahepatic biliary remnants are associated with disease severity markers and localized bile duct injury in BA. We identify T peripheral helper cells as an expanded IL-21-producing CD4⁺ T cell population within these structures, and show that post-infection IL-21 receptor blockade prevents biliary obstruction and improves survival in a murine BA model.

*How this study might affect research, practice or policy:* These findings identify the IL-21/IL-21R signaling axis as a candidate therapeutic target in BA warranting further preclinical and translational investigation. TLS maturation status in biliary remnants and serum autoantibody levels may serve as potential biomarkers of disease severity, meriting prospective evaluation in clinical cohorts.

## Introduction

Biliary atresia (BA) is a devastating neonatal liver disease characterized by progressive liver fibrosis and obliteration of extrahepatic and intrahepatic bile ducts. Although Kasai portoenterostomy restores bile flow in 50%-75% of patients, long-term outcomes remain poor, with 20-year native liver survival rates of only 27%-51%[1–4]. This complex disorder exhibits heterogeneous etiology involving both genetic susceptibility and environmental triggers, including viral infections and toxin exposure. While the initiating factors remain unclear, mounting evidence has identified immune-inflammatory dysregulation as an important contributor to BA pathogenesis[5]. Histological examination reveals marked infiltration of both innate and adaptive immune cells within the portal area of BA patients[6–11]. Of particular significance, these infiltrating lymphocytes demonstrate oligoclonal expansion within extrahepatic biliary remnants (EBR)[12,13], the pathological structures of obliterated bile ducts mixed with fibrosis tissues unique to BA patients. Supporting an autoimmune component, sera from BA patients harbor antinuclear antibodies[11] and autoantibodies targeting biliary epithelium[14,15]. Furthermore, B cell depletion significantly reduces liver damage[11], directly demonstrating the pathogenic role of humoral immunity in BA progression. Despite these advances, the specific mechanisms by which immune dysregulation drives BA pathogenesis remain poorly understood.

Ectopic lymphoid aggregates have been described in biliary remnants of BA patients, and tertiary lymphoid structures (TLSs) are increasingly recognized as critical organizers of local adaptive immune responses in chronic inflammatory and autoimmune conditions. However, the systematic characterization of TLS maturation states in BA, their relationship to disease severity, and the specific cellular mechanisms by which they may drive bile duct injury have not been elucidated. In particular, the role of T peripheral helper (Tph) cells, a specialized CD4^+^ T cell subset recently implicated in TLS-associated autoimmunity in rheumatoid arthritis, lupus, and IgG4-related disease, has not been explored in BA.

Here, we employed an integrated approach combining bulk RNA sequencing, spatial transcriptomics, single-cell RNA sequencing, multiplex immunohistochemistry, and functional validation in a rotavirus-induced murine model to characterize the immunopathological landscape of BA (Fig. 1). Our analysis reveals that TLSs within extrahepatic biliary remnants harbor expanded populations of IL-21-producing Tph cells that are associated with local autoantibody production and bile duct injury. IL-21 receptor blockade in a BA mouse model ameliorated biliary obstruction and improved survival, suggesting that IL-21/IL-21R signaling contributes to disease progression and may represent a therapeutic target warranting further investigation.

**Fig. 1:**
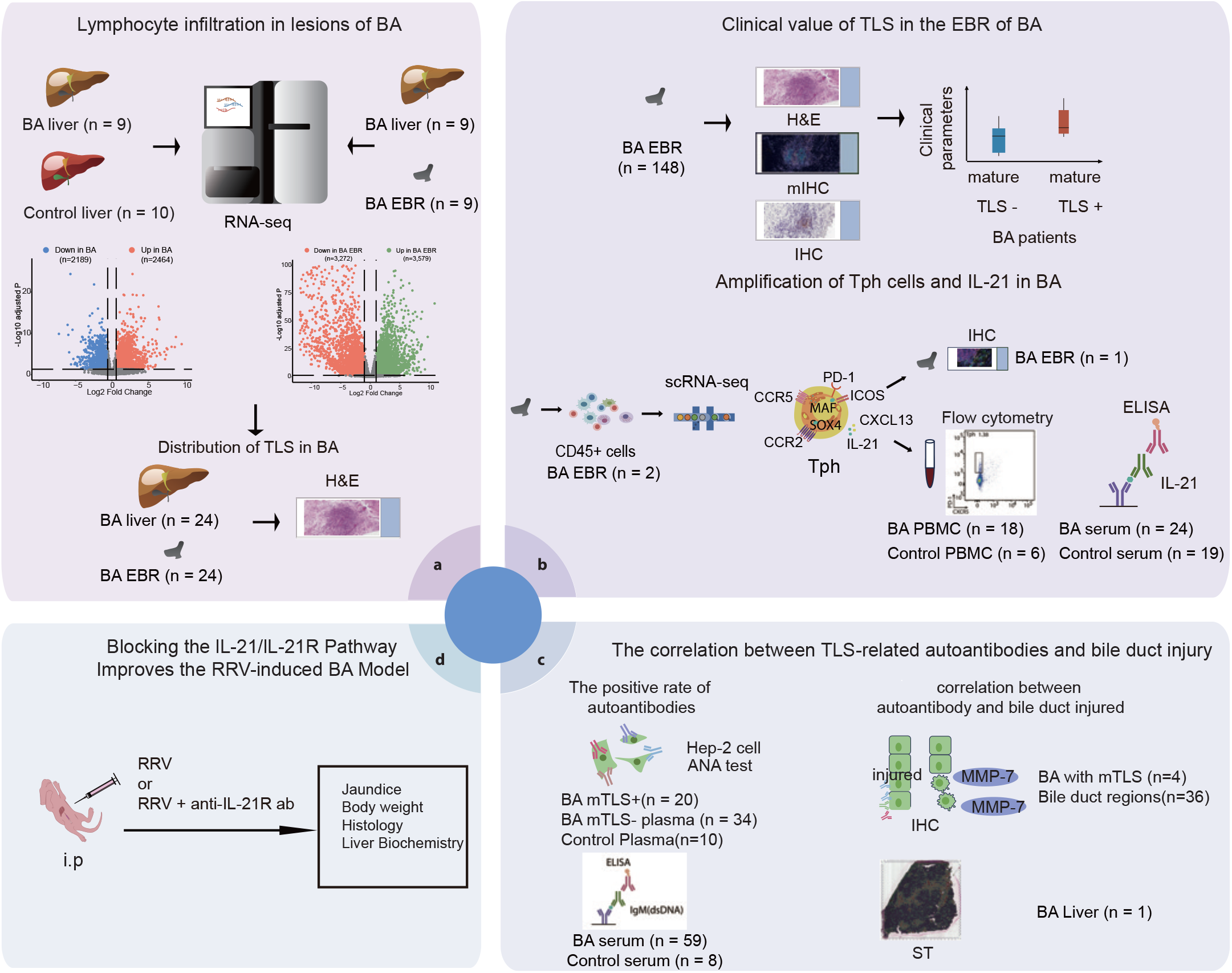
Study design and experimental workflow. Schematic overview of participant selection, experimental design, and methodological approach. BA, biliary atresia; EBR, extrahepatic biliary remnants; TLS, tertiary lymphoid structures; mTLS, mature TLS; H&E, hematoxylin and eosin staining; IHC, immunohistochemistry; mIHC, multiplex immunohistochemistry; ELISA, enzyme-linked immunosorbent assay; ANA, antinuclear antibody; RRV, rhesus rotavirus; ST, spatial transcriptomics.

## Methods

### Patients

Patients were recruited from the Departments of Pediatric Surgery at Children’s Hospital of Fudan University. Type III BA diagnosis was established through laparoscopic exploration and intraoperative cholangiography. Age-matched non-BA infantile cholestasis controls comprised children with normal liver function undergoing surgery for choledochal cysts and patients diagnosed with non-BA cholestasis using identical diagnostic procedures.

The study was approved by the Ethics Committee of Children’s Hospital of Fudan University (approval number: 2020[153]), and written informed consent was obtained from legal guardians of all participants. Patients or the public were not involved in the design, or conduct, or reporting, or dissemination plans of our research. Detailed clinical characteristics, including age at surgery and relevant biochemical measurements, are provided in Supplementary Table 4.

### RNA-seq

Total RNA was extracted from human liver specimens using TRIzol reagent (Thermo Fisher Scientific) according to the manufacturer’s protocol. RNA-seq libraries were constructed using the Fast RNA-seq Library Prep Kit V2 (ABclonal). Library quality was assessed using the Agilent 2100 Bioanalyzer (Agilent Technologies), and concentrations were quantified using a Qubit 2.0 Fluorometer (Thermo Fisher Scientific). Sequencing was performed on the Illumina NovaSeq 6000 platform generating 150 bp paired-end reads.

Raw reads were processed with TrimGalore (v0.6.10) to remove adapters and low-quality sequences. Processed reads were aligned to the human reference genome (GRCh38) using STAR (v2.5.3a)[16]. Gene expression was quantified using featureCounts (v2.0.6)[17]. Differentially expressed genes (DEGs) were identified using DESeq2 (v1.40.2)[18]: BA liver versus CC liver, |log2FoldChange| ≥ log_2_(1.5) and adjusted p-value < 0.05; BA EBR versus paired liver tissue, |log2FoldChange| ≥1 and adjusted p-value < 0.05. Volcano plots were generated using EnhancedVolcano R package (v1.18.0). Gene Set Enrichment Analysis was performed using clusterProfiler (v4.8.3)[19]. Lymphocyte infiltration levels were estimated using TIMER2.0, which employs a computational algorithm to calculate the sum of CD4^+^ T cell, CD8^+^ T cell, and B cell infiltration scores[20]. TLS signature scores were calculated using a 7-gene signature[21] with the ssGSEA algorithm from the GSVA package[22].

### Evaluation and quantification of ectopic lymphoid aggregates (ELAs)

Liver and EBR specimens from BA patients were collected during surgery, formalin-fixed, paraffin-embedded, and sectioned at 5.0 μm thickness. All EBR specimens were obtained from the hilar region during Kasai portoenterostomy. Sections were stained with hematoxylin and eosin (H&E) for morphological evaluation. ELAs were identified as discrete lymphocyte aggregates and quantified using QuPath software (v0.5.1) according to previously established criteria[23,24]. Representative images of ELAs in both liver and EBR tissues are shown in Fig. 2h.

**Fig. 2:**
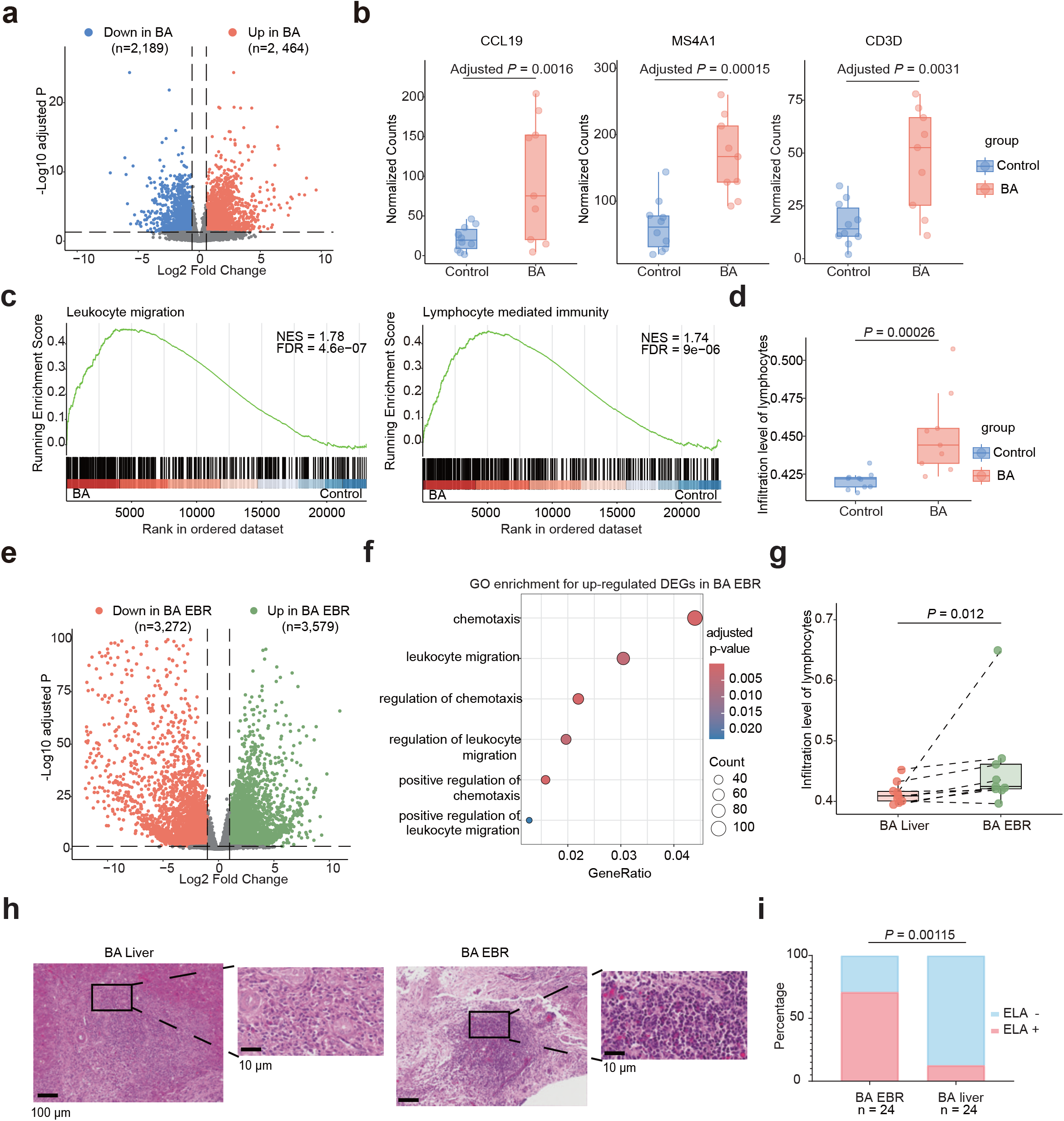
Immune activation and ectopic lymphoid aggregate formation in biliary atresia. (a) Volcano plot showing differentially expressed genes between BA (n = 9) and age-matched choledochal cyst control (n = 10) liver tissues. A total of 2,464 genes were upregulated (red) and 2,189 genes were downregulated (blue) in BA liver (criteria: |log2fold change| ≥ log_2_(1.5) and adjusted *P* < 0.05). (**b**) Box plots showing mRNA expression levels of *CCL19*, *MS4A1*, and *CD3D* in BA liver (n = 9) versus control liver tissues (n = 10). Adjusted *P*-values are shown at the top. (**c**) Gene Set Enrichment Analysis (GSEA) of indicated pathways comparing BA liver with control liver tissues. FDR, false discovery rate; NES, normalized enrichment score. (**d**) Differential lymphocyte infiltration levels estimated using TIMER2 (Mann–Whitney U test). (**e**) Volcano plot showing differentially expressed genes between BA extrahepatic biliary remnants (EBRs) and paired liver tissues (n = 9). A total of 3,579 genes were upregulated (green) and 3,272 genes were downregulated (red) in BA EBRs (criteria: |log2fold change| ≥ 1 and adjusted *P* < 0.05). (**f**) Gene Ontology (GO) enrichment analysis of upregulated differentially expressed genes (DEGs) in BA EBRs, highlighting leukocyte-related biological processes. (**g**) Lymphocyte infiltration levels in BA EBRs versus paired liver tissues (n = 9), estimated using TIMER2 (Wilcoxon signed-rank test). (**h**) Representative hematoxylin and eosin (H&E) images showing ectopic lymphoid aggregates (ELAs) in BA EBR and liver tissues. (**i**) Quantification of ELA prevalence in paired EBR and liver tissues from BA patients (n = 24) (McNemar’s test).

### Multiplex immunohistochemical staining

Multiplex immunohistochemical staining was performed using the mIHC Fluorescence kit (Recordbio Biological Technology) according to the manufacturer’s protocol. Paraffin-embedded sections were deparaffinized in xylene, rehydrated, and subjected to antigen retrieval via microwave heating for 15 minutes. After cooling, endogenous peroxidase activity was blocked using 3% H₂O₂ for 20 minutes in darkness. Sections were then blocked with 10% goat serum for 1 hour to prevent non-specific binding. Primary antibodies were applied sequentially and incubated overnight at 4°C: anti-CD20 (Abcam, ab78237), anti-CD3 (Abcam, ab237707), anti-CD4 (Abcam, ab133616), anti-PD1 (Abcam, ab237728), anti-CXCR5 (Abcam, ab254415). Following primary antibody incubation, sections were incubated with HRP-conjugated secondary antibody for 1 hour, followed by tyramide signal amplification (TSA) with fluorophores. Microwave heat treatment for 15 minutes was performed after each TSA step to remove antibodies before subsequent staining. After completing all antigen labelling, nuclei were counterstained with DAPI. Stained slides were digitized and analyzed using KFBIO Digital Pathology Slide Scanners.

### Immunohistochemistry

Following rehydration, paraffin-embedded tissue sections underwent antigen retrieval by microwave heating for 15 minutes. After cooling, endogenous peroxidase activity was blocked using 3% H₂O₂ for 20 minutes. Sections were then blocked with QuickBlock™ Blocking Buffer for Immunol Staining (Beyotime) for 20 minutes. Primary antibodies were applied overnight at 4°C: anti-CD23 (Long Island, M-0104), anti-MMP-7 (Abcam, ab207299), and anti-CK19 (Abcam, ab52625). Following primary antibody incubation, sections were incubated with appropriate HRP-conjugated secondary antibodies. Immunoreactivity was visualized using 3,3’-diaminobenzidine (DAB) substrate (Beyotime), followed by hematoxylin counterstaining. Images were acquired and analyzed using an Olympus VS 200 scanner. For immunoglobulin deposition analysis, consecutive sections were stained with an HRP-conjugated goat anti-human IgG antibody (Abcam, ab98624) and an HRP-conjugated goat anti-human IgM antibody (Abcam, ab97205).

### TLS assessment

TLSs were defined as organized aggregates of infiltrating T and B lymphocytes[25]. Cellular composition within TLS was determined using CD3 and CD20 immunofluorescence staining to identify T and B cell populations, respectively. TLS maturity was assessed by CD23 immunohistochemistry[25], where sections were categorized as mTLS+ if at least one CD23^+^ follicular dendritic cell was detected within any TLS structure, indicating the presence of a nascent or established germinal center. Sections lacking CD23^+^ signals were classified as mTLS-.

### Correlation analysis between TLS maturity and MMP-7 expression

The correlation between TLS maturity and MMP-7 expression levels in bile ducts was assessed using immunohistochemistry. CK19 immunostaining was performed to identify bile ducts. MMP-7 expression was quantified in bile duct regions located within 1 mm of TLS structures on adjacent consecutive sections. Positive areas, defined as diaminobenzidine (DAB)-positive brown staining, were quantified automatically using QuPath software. The MMP-7 expression index was calculated as:

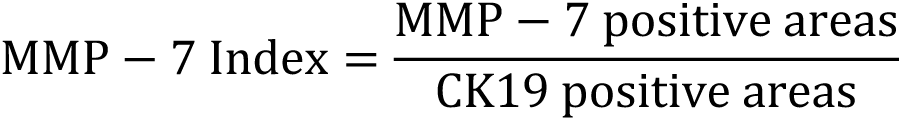

### Correlation analysis between TLS maturity and hepatic fibrosis

The correlation between TLS maturity and hepatic fibrosis was assessed using liver tissue sections from BA patients. Hepatic fibrosis was evaluated by experienced pathologists according to the Batts-Ludwig scoring system[26].

### Correlation analysis of MMP-7 expression and immunoglobulin deposition

To investigate the relationship between MMP-7 expression and immunoglobulin deposition in bile ducts, we performed analysis on extrahepatic biliary remnant sections from 4 patients with mature TLS. For each patient, 4 consecutive sections were prepared. Section 1 was stained with CK19 to identify bile ducts. Section 2 was stained for MMP-7. Sections 3 was stained with an HRP-conjugated goat anti-human IgG antibody (Abcam, ab98624), and section 4 was stained with an HRP-conjugated goat anti-human IgM antibody (Abcam, ab97205). Regions of interest (ROIs) were defined based on CK19^+^ bile duct areas from section 1 and applied to corresponding areas in sections 2-4. MMP-7- and IgG-positive areas within ROIs were quantified automatically using QuPath software, yielding MMP-7/ROI and IgG/ROI ratios for each bile duct.

### Single-cell RNA sequencing

For single-cell RNA sequencing, extrahepatic biliary remnants from two BA patients were combined, mechanically fragmented and enzymatically digested with DNase I (Sigma-Aldrich) and collagenase IV (Gibco) at 37°C with agitation for 30 minutes. Following digestion, cell suspensions were filtered through 70-μm cell strainers, washed with PBS, and centrifuged. Cells were labeled with CD45 magnetic microbeads (Miltenyi Biotec) for 15 minutes at 4°C and positively selected using magnetic separation. Single-cell RNA sequencing library was constructed from isolated CD45^+^ cells using the 10x Genomics Chromium system and sequenced on the Illumina NovaSeq 6000 platform.

Raw sequencing data were converted to FASTQ files using Illumina bcl2fastq (v2.19.1) and aligned to the human genome reference (GRCh38) using the CellRanger (v7.1.0) to generate feature-barcode matrix. Data analysis was performed using Seurat (v4.4.0)[27]. Quality control retained cells with 200-6,000 detected genes and mitochondrial gene percentage below 10%. Doublets were identified and removed using DoubletFinder[28]. After quality control, 19,858 genes × 10,821 cells were retained for analysis. Data normalization and scaling were performed using SCTransform. Principal component analysis (PCA) was conducted for dimensionality reduction, with significant principal components determined using ElbowPlot. These components were used to construct shared nearest neighbor (SNN) graphs using FindNeighbors and perform clustering using FindClusters. Uniform Manifold Approximation and Projection (UMAP) was applied for two-dimensional visualization. Differentially expressed genes for each cluster were identified using FindAllMarkers, and cell clusters were annotated based on canonical lineage markers. Ligand-receptor interaction analysis was performed using CellChat[29,30].

### ELISA

Serum concentrations of IL-21 and anti-dsDNA IgM were measured using ELISA kits (Enzyme-linked Biotechnology) according to the manufacturer’s protocol. Briefly, diluted serum samples (1:5 dilution) and standards were added to 96-well plates pre-coated with specific capture antibodies. Following incubation, biotinylated detection antibodies were added. After thorough washing, streptavidin-HRP conjugate was added. Color development was initiated by adding TMB substrate. Optical density was measured at 450 nm using a microplate spectrophotometer, and analyte concentrations were determined from standard curves.

### In vitro PBMC stimulation and intracellular cytokine staining

PBMCs were isolated and stimulated with the leukocyte activation cocktail (BD Biosciences, 568685) at a density of 2-3 × 10^6^ cells/mL in RPMI 1640 medium (GIBCO) for 5 hours at 37°C under 5% CO_2_, with unstimulated cells serving as negative controls. Following stimulation, cells were stained with Fixable Viability Dye 780 (STARTER, S0D0026) and then incubated with a surface antibody cocktail comprising antibodies against CD3 (BD Pharmingen, S0B5722), CD4 (570234), CD8a (568685), PD-1 (562516), CXCR5 (563105), and CD45 (560976). After surface staining, cells were fixed, permeabilized, and stained intracellularly for IL-17A (563746) and IL-21 (562042). All samples were washed, resuspended in dilution buffer, filtered through a 70 μm strainer, and analyzed on a BD Fortessa flow cytometer using FACSDiva software. Data analysis was performed with FlowJo (v10.0.7).

### Flow cytometry

PBMCs were isolated using Ficoll-Paque PLUS (Cytiva) and resuspended in FACS buffer containing PBS supplemented with 1% FBS and 1× penicillin-streptomycin. All staining procedures were performed in darkness to preserve fluorochrome integrity. Cell viability was assessed using Zombie Yellow (BioLegend) for 30 minutes in PBS. Following washing, cells were incubated with fluorochrome-conjugated antibodies against surface markers for 15 minutes at room temperature in FACS buffer: BUV395 anti-Human CD3 (BD Biosciences, 564001), Brilliant Violet 510 anti-human CD4 (BioLegend, 300546), Alexa Fluor 700 anti-human CD8a (BD Biosciences, 557945), Brilliant Violet 421 anti-human PD-1 (BioLegend, 329920), and Alexa Fluor 488 anti-human CXCR5 (BD Biosciences, 558112). Cells were washed with cold PBS, filtered through 70-μm mesh, and analyzed using a BD Fortessa flow cytometer with FACSDiva software. Data analysis was performed using FlowJo (v10.0.7). For intracellular IL-21 staining, PBMCs were stimulated with PMA and ionomycin in the presence of brefeldin A, followed by fixation, permeabilization, and staining with an anti-IL-21 antibody.

### Antinuclear antibody (ANA) analysis

ANA detection and pattern assessment were performed using HEp-2 cells as substrate. HEp-2 cells were seeded in 24-well plates at 2 × 10^5^ cells per well and cultured for 24 hours. Cells were then fixed with 4% paraformaldehyde (Servicebio) for 10 minutes, permeabilized with 1% Triton X-100 (Sangon Biotech) for 10 minutes, and blocked with 2% bovine serum albumin (Sangon Biotech) for 1 hour at room temperature. Cells were incubated with diluted patient serum samples (1:100) in blocking buffer overnight at 4°C. Following primary incubation, cells were washed and stained with Alexa Fluor 488-conjugated goat anti-human IgG (Sangon Biotech, D110164) for 1 hour at room temperature. Fluorescence imaging and ANA pattern analysis were performed using a Leica DMi8 microscope.

### Establishment and analysis of the BA mouse model

All animal experiments were conducted in accordance with the National Institutes of Health Guide for the Care and Use of Laboratory Animals and were approved by Animal Care and Use Committee of School of Basic Medical Sciences, Fudan University with the reference number 20241213-002. Pregnant BALB/c mice at 16 days of gestation were purchased from Biocytogen and housed in SPF-level facilities at the Laboratory Animal Center, Fudan University (temperature 20-25°C, humidity 40%-70%, 12-hour light-dark cycle, with free access to food and water). Delivery status was monitored daily at 9:00 and 16:00. The day of birth was designated as day 0. Newborn BALB/c mice were intraperitoneally injected with 25 μl of rhesus rotavirus (RRV; 1.5×10^6^ pfu/ml) within 12-24 hours after birth to establish the experimental BA model. Control mice received an equal volume of serum-free Dulbecco’s Modified Eagle Medium (DMEM, GIBCO) containing 0.5 μg/ml trypsin (RRV culture medium).

For IL-21/IL-21R pathway blockade, neonatal mice from the same litter were randomly allocated to the anti-IL-21R antibody treatment or untreated RRV-induced group on day 0. Mice received 10 μg of anti-IL-21R antibody (Bio X Cell, BE0258) per mouse by intraperitoneal injection beginning on day 4 after RRV administration, followed by additional injections (10 μg/mouse) every 48 hours through day 10. Mice dying within 2 days of birth were excluded from analysis. Throughout the experiment, weight gain, jaundice (assessed by skin color in hairless areas), and stool/urine color were monitored and compared between groups. Animals were sacrificed under anesthesia on day 12.

Liver samples were processed into paraffin sections for H&E staining. Extrahepatic bile duct tissues were fixed in 4% paraformaldehyde, cryoprotected in 15% and 30% sucrose solutions, embedded in OCT compound, and sectioned at 10 μm thickness for H&E staining. Serum samples were collected on day 12 and analyzed for levels of γ-glutamyl transpeptidase (GGT), direct bilirubin (DBIL), alanine aminotransferase (ALT), and aspartate aminotransferase (AST) using a BS-240 - Clinical Chemistry Analyzer (Mindray).

### Statistical analyses

Data analysis and visualization for bulk RNA sequencing and single-cell RNA sequencing (scRNA-seq) were performed using R (v4.3.0). GraphPad Prism 9 was used for statistical analysis and graphical representation of experimental data. Data normality was evaluated using the Shapiro-Wilk test. For normally distributed continuous variables, groups were compared using the Student’s t-test. For non-normally distributed data, the Mann-Whitney U test or Wilcoxon signed-rank test was applied. Categorical variables were compared using Fisher’s exact test. Correlations were assessed using Spearman’s rank correlation (for non-normally distributed data) or Pearson’s correlation (for normally distributed data). Statistical significance was defined as *P* < 0.05.

## Results

### Immune dysregulation and ectopic lymphoid aggregate formation in biliary atresia lesions

To systematically characterize the immune microenvironment underlying BA pathogenesis, we performed RNA-seq analysis on liver biopsies from 9 BA patients and 10 age-matched choledochal cyst controls. We identified 4,653 differentially expressed genes (Fig. 2a), with BA livers demonstrating significant upregulation of genes involved in lymphocyte chemotaxis (*CCL19*, *CXCL10*, *CCL28*, *CXCL9*), B cell activation (*MS4A1*, *IGHG2*, *IGHA1*, *IGHD*), and T cell function (*CD3D*, *CD8A*, *GZMA*) (Fig. 2b and Supplementary Fig. 1a). These findings are consistent with prior studies that have documented immune cell infiltration and B cell tolerance defects in BA livers[7,11]. Gene set enrichment analysis (GSEA) confirmed elevated expression of immune-related pathways in BA livers, including leukocyte migration and lymphocyte-mediated immunity (Fig. 2c), along with hallmark gene sets for inflammatory responses (Supplementary Fig. 1b). Computational deconvolution using TIMER2.0, an algorithm to estimate infiltration levels of lymphocytes from RNA-seq data[20], revealed significantly increased lymphocyte infiltration in BA livers compared to controls (*P* = 0.00026, Mann-Whitney U test) (Fig. 2d).

Having established the inflammatory nature of BA liver, we next investigated extrahepatic biliary remnants (EBRs), the pathological structures of obliterated bile ducts outside the liver that represent a distinctive lesion site unique to BA patients. To characterize the immune landscape of these structures, we performed RNA-seq analysis on paired EBR and liver tissues from nine BA patients (Fig. 2e). Gene Ontology (GO) analysis revealed that EBRs exhibited significant enrichment of pathways associated with lymphocyte chemotaxis and migration compared to matched liver tissues (Fig. 2f). Consistent with this observation, computational deconvolution demonstrated significantly higher lymphocyte infiltration in EBRs than in liver tissues from the same patients (*P* = 0.012, Wilcoxon signed-rank test) (Fig. 2g).

To validate these transcriptomic findings and evaluate lymphocyte distribution patterns across BA lesion sites, we performed hematoxylin and eosin (H&E) staining on paired liver and EBR tissues from an independent cohort of 24 BA patients. We observed that lymphocytes frequently formed ectopic lymphoid aggregates (ELAs) in both tissue types (Fig. 2h), with significantly higher prevalence in EBRs than in livers (*P* = 0.00115, McNemar’s test) (Fig. 2i). The formation of lymphoid aggregates in biliary remnants is consistent with earlier histopathological observations in BA[31]. These ELAs morphologically resembled tertiary lymphoid structures (TLSs)[23,24]. To confirm their identity as bona fide TLSs, we performed multiplex immunohistochemistry (mIHC) staining, which revealed that ELAs in EBR tissues were predominantly composed of CD20^+^ B cells and CD3^+^ T cells (Fig. 3a). Supporting this classification, TLS signature analysis[21] of RNA-seq data revealed a significantly elevated TLS signature in EBR tissues compared to paired liver samples (*P* = 0.044, paired Student’s *t* test) (Supplementary Fig. 1c). These findings define a distinct inflammatory landscape in BA characterized by marked lymphocyte infiltration and TLS formation, with EBRs representing preferential sites of ectopic lymphoid neogenesis.

**Fig. 3:**
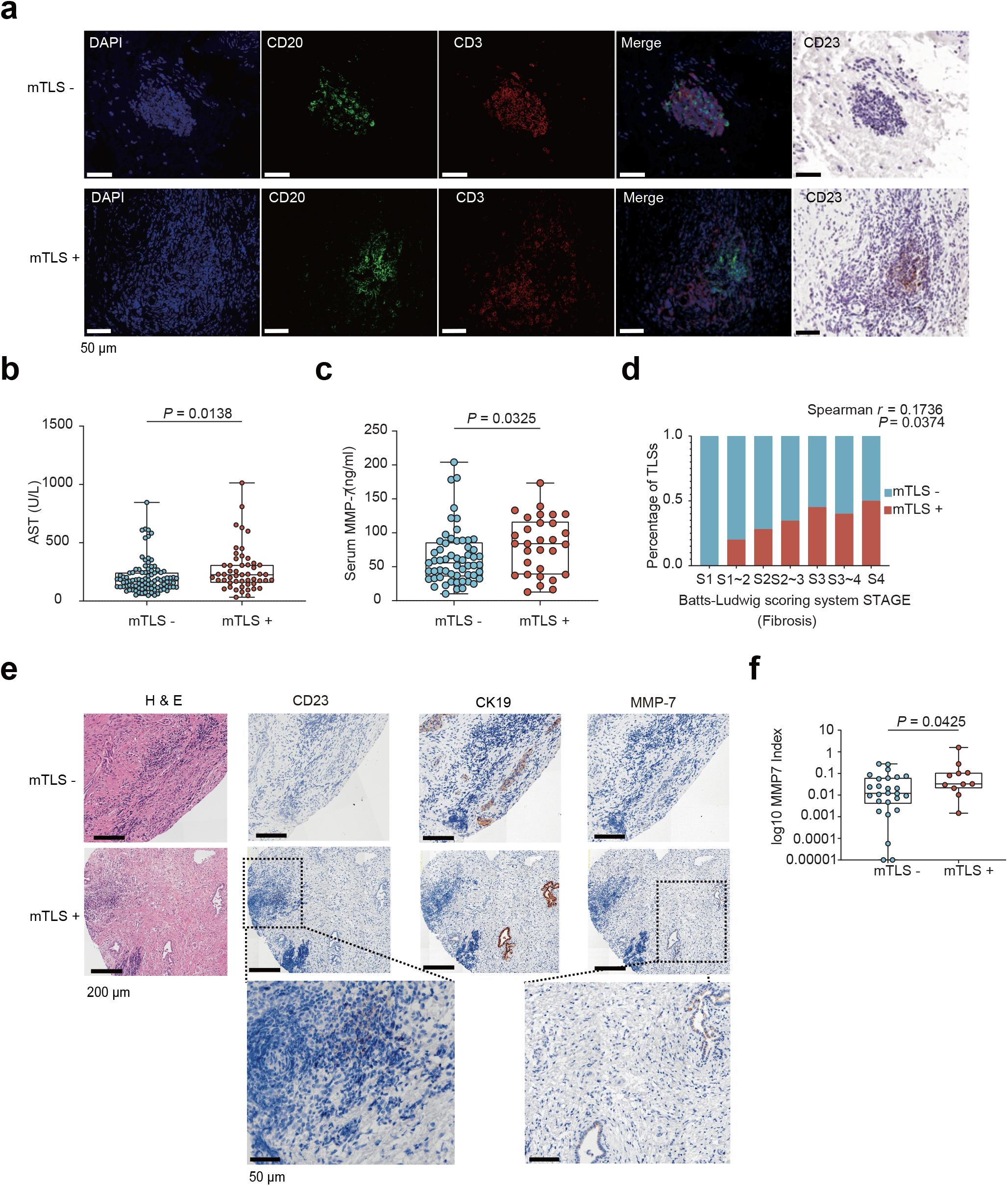
Mature tertiary lymphoid structures in EBRs associate with disease severity markers. (a) Identification of tertiary lymphoid structures (TLSs) in EBRs from BA patients by multiplex immunohistochemistry for CD3 (red) and CD20 (green). TLS maturation was assessed by CD23 immunohistochemistry (brown) on consecutive sections. mTLS+, patients with mature TLS; mTLS-, patients without mature TLS. (**b**) Serum aspartate aminotransferase (AST) levels in BA patients stratified by mature TLS status (mTLS-, n = 95; mTLS+, n = 49) (Mann-Whitney U test). (**c**) Serum matrix metalloproteinase-7 (MMP-7) levels in BA patients stratified by mature TLS status (mTLS-, n = 57; mTLS+, n = 30) (Mann-Whitney U test). (**d**) Correlation between Batts-Ludwig fibrosis stage and mature TLS presence in EBRs from BA patients. y-axis “Percentage of TLSs” denotes the proportion of mTLS+ (red) and mTLS− (blue) patients per stage. *r*, Spearman correlation coefficient. (**e**) Representative consecutive tissue sections showing TLS maturation and MMP-7 expression in CK19^+^ EBRs. From Left: H&E staining showing TLS morphology; CD23 staining indicating TLS maturation; MMP-7 expression in CK19^+^ biliary epithelium. (**f**) MMP-7 index in BA patients stratified by mature TLS status (mTLS-, n = 26; mTLS+, n = 11) (Mann-Whitney U test).

### Mature TLSs in BA EBRs associate with disease severity

Like secondary lymphoid organs, TLSs can be classified as mature when they contain CD23^+^ germinal centers, where organized cell-cell interactions occur to generate robust adaptive immune responses[32]. To assess the clinical significance of TLS maturation in BA, we performed CD20 and CD3 mIHC staining on EBR tissues from 148 BA patients, followed by CD23 immunohistochemistry to classify patients based on TLS maturation state (Fig. 3a). The classification of mature TLS (mTLS+) was defined as the presence of at least one CD23^+^ follicular dendritic cell within any TLS structure, consistent with established criteria for identifying germinal center formation in ectopic lymphoid tissues[25]. Patients with mTLS+ demonstrated more severe disease manifestations compared to those without mature TLS (mTLS-). Specifically, the mTLS+ group had significantly elevated serum aspartate aminotransferase (AST) levels, a marker of hepatocellular injury (*P* = 0.0138, Mann-Whitney U test) (Fig. 3b). Matrix metalloproteinase-7 (MMP-7), a key enzyme in tissue remodeling associated with liver damage and fibrosis progression in BA[33,34], was also significantly higher in the mTLS+ group (*P* =0.0325, Mann-Whitney U test) (Fig. 3c). Furthermore, liver fibrosis stage, assessed by the Batts-Ludwig scoring system, showed a modest but statistically significant positive correlation with the percentage of mTLS+ patients (Spearman *r* = 0.1736; *P* = 0.0374) (Fig. 3d).

To examine whether mature TLSs are spatially associated with local tissue damage, we assessed bile duct injury in their vicinity. Since MMP-7 is primarily released by injured cholangiocytes[33,34] and serves as a marker of bile duct damage, we quantified MMP-7 expression in cholangiocytes within 1 mm of TLSs. Using immunohistochemistry, we measured the MMP-7 positive area within CK19^+^ bile ducts and found significantly higher MMP-7 expression in sections containing mature TLSs compared to those without (*P* = 0.0425, Mann-Whitney U test) (Fig. 3e, f), indicating that mature TLSs may contribute to local bile duct injury.

### Identification of disease-associated Tph cells in BA

To investigate the cellular mechanisms underlying TLS formation and function in BA EBRs, we performed single-cell RNA-seq (scRNA-seq) on CD45^+^ cells isolated from EBR tissues with mature TLSs (Fig. 4a). After preprocessing and quality control, a total of 10,821 cells were partitioned into 17 clusters (Fig. 4b) based on canonical marker gene expression (Supplementary Fig. 2a and Supplementary Table 1). Given the critical role of CD4^+^ T cells in TLS formation and function[35,36], we further partitioned the 1,424 identified CD4^+^ T cells into nine distinct subsets (Fig. 4c, Supplementary Fig. 2b and Supplementary Table 2). Among them, we identified a subset of T peripheral helper (Tph) cells based on the absence of the follicular helper marker *CXCR5* alongside high expression of *CXCL13*, *IL21*, and *PDCD1* (encoding *PD-1*) (Fig. 4d and Supplementary Fig. 2b), consistent with their initial description in rheumatoid arthritis synovium[37]. These cells, typically located at TLS peripheries, orchestrate lymphoid organization by recruiting B and T cells through CXCL13 secretion and promoting B cell differentiation via IL-21 in autoimmune disease pathogenesis[38].

**Fig. 4:**
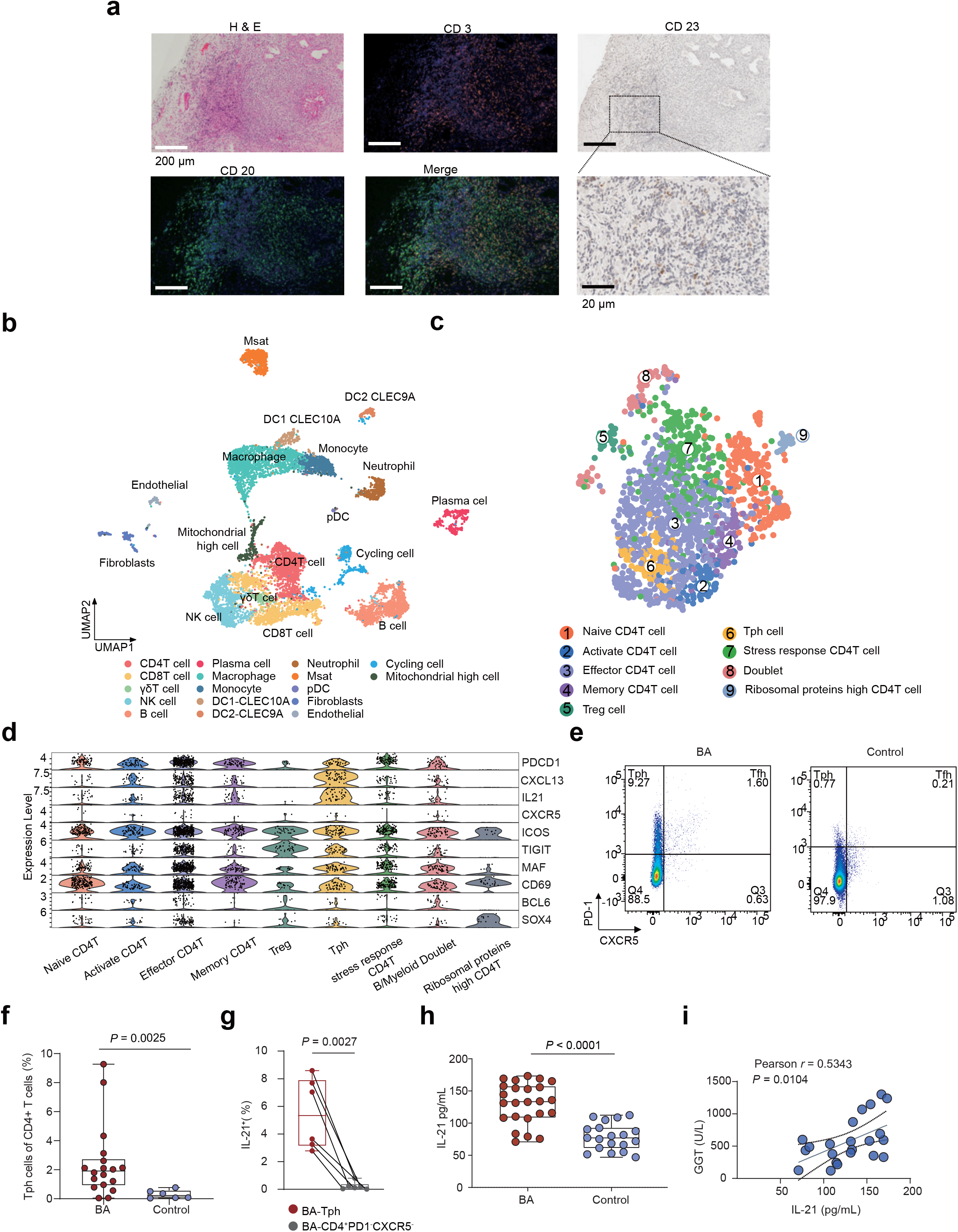
T peripheral helper cells are expanded in BA compared with age-matched controls. (**a**) H&E staining and multiplex immunohistochemistry for CD3 (red), CD20 (green), and CD23 (brown) in a representative TLS from BA EBR tissue used for scRNA-seq. (**b**) Uniform Manifold Approximation and Projection (UMAP) visualization of all cells with cluster annotations. (**c**) UMAP visualization of CD4^+^ T cells with cluster annotations. (**d**) Violin plots showing expression of marker genes for each CD4^+^ T cluster. (**e**) Representative flow cytometry plots showing T peripheral helper (Tph) cells (CD3^+^CD4^+^CD8a^-^PD-1^hi^CXCR5^-^) in peripheral blood mononuclear cells (PBMCs) from a patient with BA and a control. (**f**) Frequency of Tph cells in PBMCs from BA patients (n = 18) and age-matched controls with non-BA infantile cholestasis (n = 6) (Mann-Whitney U test). (**g**) Frequencies of IL-21^+^ cells in Tph cells vs. CD3^+^CD4^+^CD8a^-^PD-1^-^CXCR5^-^ countparts from stimulated PBMCs of BA patients (n = 6). (**h**) Serum interleukin-21 (IL-21) concentrations in BA patients (n = 24) and age-matched controls with non-BA infantile cholestasis (n = 19) (Mann-Whitney U test). (**i**) Correlation between serum IL-21 and γ-glutamyl transferase (GGT) levels in patients with BA (n = 21) (Pearson correlation analysis).

Having identified Tph cells within EBR tissues, we next examined their presence in the peripheral blood. Flow cytometric analysis (Supplementary Fig. 3a) revealed significantly increased frequencies of CD4^+^PD-1^hi^CXCR5^-^ Tph cells in BA patients compared to age-matched non-BA infantile cholestasis controls (*P* = 0.0025, Mann-Whitney U test) (Fig. 4e, f). The frequency of circulating Tph cells showed positive trends towards correlation with serum AST (Spearman *r* = 0.37, *P* = 0.0757) and ALP levels (Spearman *r* = 0.35, *P* = 0.0975) (Supplementary Fig. 3b, c), suggesting that Tph-mediated immunity may contribute to hepatic injury progression.

To investigate whether circulating Tph cells are capable of producing IL-21, a key cytokine that drives B cell differentiation[38], we stimulated PBMCs from BA patients with PMA and ionomycin and performed intracellular IL-21 staining. Tph cells expressed significantly higher levels of IL-21 compared to their CD4⁺PD-1⁻CXCR5⁻ counterparts (Fig. 4g and Supplementary Fig. 3f, g). Notably, circulating CD4⁺PD-1^hi^CXCR5⁺ T follicular helper (Tfh) cells, another CD4⁺ T cell subset capable of IL-21 production that shares substantial functional overlap with Tph cells[38], were present at extremely low frequencies under these conditions, precluding reliable assessment of their IL-21 producing capacity (Supplementary Fig. 3d, e). Together, these results confirm the capacity of Tph cells to produce IL-21 ex vivo.

We next measured serum IL-21 levels by enzyme-linked immunosorbent assay (ELISA). BA patients exhibited significantly elevated IL-21 levels compared to controls (*P* <0.0001, Mann-Whitney U test) (Fig. 4h). Furthermore, serum IL-21 levels positively correlated with γ-glutamyl transferase (GGT) levels (Pearson *r* = 0.5343, *P* = 0.0104) (Fig. 4i), suggesting a link between systemic IL-21 and disease severity. Collectively, these findings support the expansion of Tph cells in BA and their capacity to produce IL-21, although the precise causal relationship between Tph-derived IL-21 and disease progression warrants further functional investigation.

### Tph-B cell interactions in EBRs

Having identified Tph cells as a disease-associated cell type, we next characterized the complete B cell landscape within BA EBRs to understand the potential downstream effects of Tph cell activation. Using unsupervised clustering, we identified ten B cell subsets within EBR infiltrates, including precursor B cells (pre-B cells; *CD19*^+^*RAG1*^+^*RAG2*^+^*PAX5*^+^*SOX4*^+^*IGLL1*^+^), naïve B cells (*IGHD*^+^*FCER2*^+^*SELL*^+^), activated B cells (*CD69*^+^*NR4A2*^+^*CD83*^+^*BTG1*^+^*JUND*^+^), APC-enriched B cells (*HLA-DPB1*^+^*HLA-DRB5^+^HLA-DRA*^+^), germinal center B cells (GC B cells; *BCL6^+^AICDA^+^EZH2^+^MIR155HG^+^BCL2A1^+^*), atypical memory B cells (AtM B cells; *CR2*^-^*CD27*^-^*IgD*^-^*DUSP4*^+^*ITGAX*^+^*FCRL5*^+^*ZEB2*^+^*FGR*^+^*FCRL4*^+^), plasma cells (*CD38*^+^*JCHAIN*^+^*MZB1*^+^), and plasmablasts (*MKI67*^+^) (Fig. 5a, Supplementary Fig. 4a and Supplementary Table 3).

**Fig. 5:**
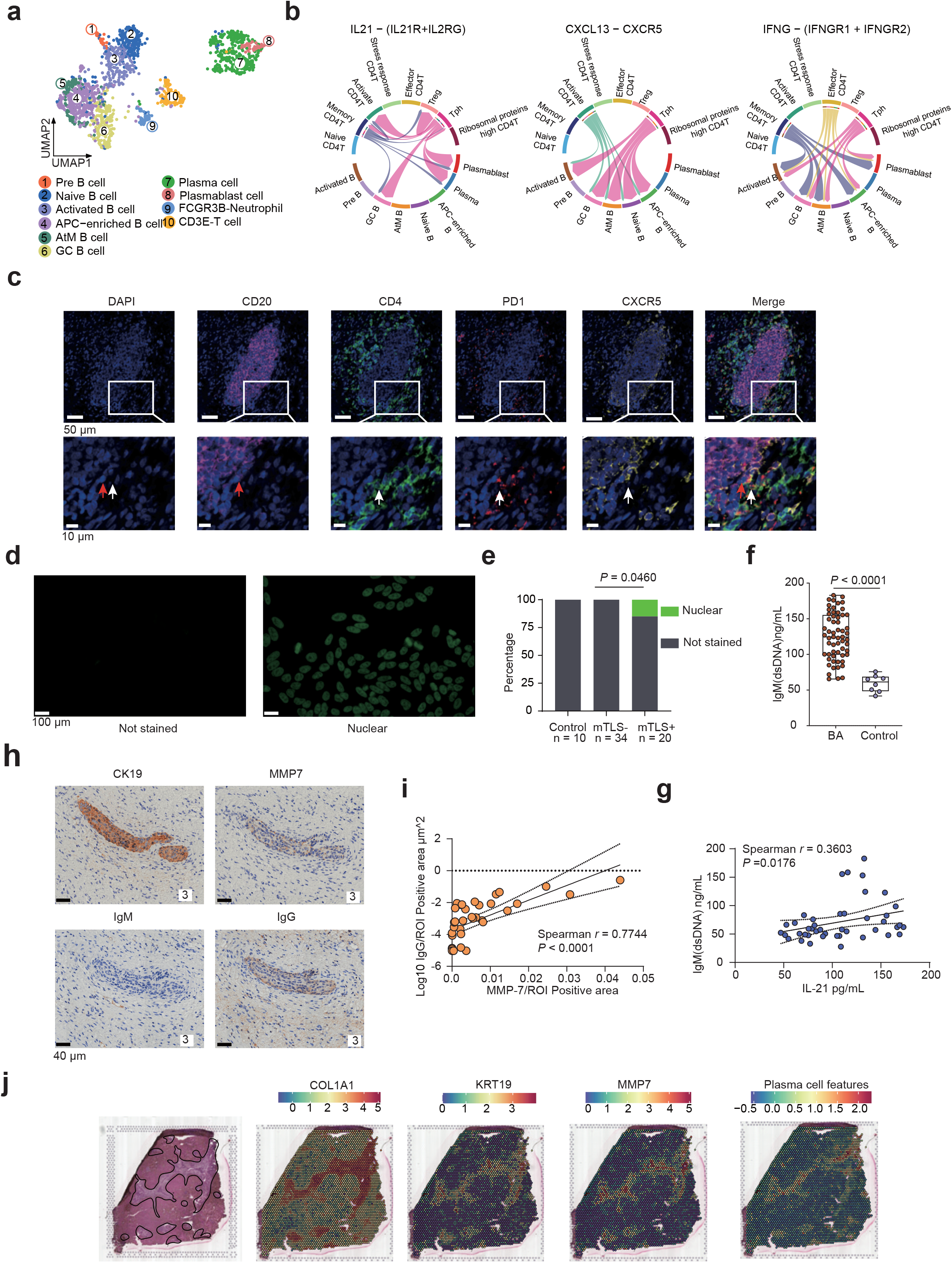
Tph-B cell interactions and autoimmune responses. (**a**) UMAP visualization of B and plasma cells with cluster annotations. (**b**) Receptor-ligand interactions (*IL-21*/*IL-21R*, *CXCL13*/*CXCR5*, and *IFNG*/*IFNGR*) between CD4^+^ T and B cell subsets. (**c**) Multiplex immunohistochemical staining showing B cells (CD20, purple) and Tph cells (CD4, green; PD-1, red; CXCR5, yellow) within TLSs. Red arrows indicate B cells (CD20^+^); white arrows indicate Tph cells (CD4^+^PD-1^+^CXCR5^-^). (**d**) Representative images showing antinuclear antibody (ANA) staining from plasma of a BA patient and a control. (**e**) ANA positivity prevalence in age-matched controls (n = 10) and BA patients stratified by mature TLS status (mTLS-, n = 34; mTLS+, n = 20) (Fisher’s exact test). (**f**) Serum anti-dsDNA IgM levels in BA patients (n = 59) and non-BA infantile cholestasis controls (n = 8) (Mann-Whitney U test). (**g**) Correlation between serum IL-21 and anti-dsDNA IgM levels in BA patients (Spearman correlation analysis). (**h**) Representative immunohistochemical images of consecutive sections showing CK19, MMP-7, IgM, and IgG expression in EBR tissue from a BA patient. (**i**) Correlation between MMP-7 and IgG density per region of interest (ROI) within the same CK19^+^ region (Spearman correlation analysis). (**j**) Spatial transcriptomic analysis showing expression of fibrosis marker (*COL1A1*), bile duct marker (*KRT19*), *MMP7*, and plasma cell gene signature (*MZB1*, *JCHAIN*, *XBP1*, *IGHA1*, *IGHG1*, *IGHG2*, *IGHG3*, *IGHG4*)[43] in BA liver.

To investigate potential T-B cell interactions in BA EBRs, we analyzed intercellular communication between B and CD4^+^ T cell subsets using our scRNA-seq data. Ligand-receptor analysis revealed that Tph cells were a prominent source of key factors associated with B cell migration and terminal differentiation through IL-21 (via IL-21R), CXCL13 (via CXCR5), and IFN-γ (via IFNGR1/2) signaling (Fig. 5b and Supplementary Fig. 4b). To validate these predicted interactions in situ, we performed multiplex immunofluorescence staining for CD4^+^PD-1^hi^CXCR5^-^ Tph cells and CD20^+^ B cells. Our result revealed that Tph cells localized predominantly at TLS peripheries in close proximity to B cells (Fig. 5c), consistent with a role in orchestrating B cell recruitment and differentiation within EBR-associated TLSs.

### Autoimmune responses in TLSs are associated with biliary injury

In autoimmune diseases, TLSs at inflammatory sites respond to local antigens and are associated with autoantibody production[39–41]. Given the presence of mature TLSs in BA EBRs and their association with disease severity, we investigated whether these structures similarly associate with autoantibody production. Antinuclear antibody (ANA) staining (Fig. 5d) revealed higher ANA positivity in the mTLS+ group compared to mTLS-patients (*P* = 0.0460, Fisher’s exact test) (Fig. 5e), suggesting a link between mature TLSs and systemic autoantibody production in BA. We next quantified serum anti-dsDNA IgM, a key systemic lupus erythematosus marker known to mediate tissue injury[42]. Serum anti-dsDNA IgM was significantly elevated in BA patients compared to age-matched non-BA cholestasis controls (*P* < 0.0001, Mann-Whitney U test) (Fig. 5f). Furthermore, we observed a significant positive correlation between serum IL-21 and anti-dsDNA IgM levels (Spearman *r* = 0.3603, *P* = 0.0176) (Fig. 5g), indicating that IL-21 potentially drives TLS-mediated autoantibody production in BA.

To explore the spatial relationship between autoantibody deposition and bile duct damage in BA, we examined immunoglobulin deposition and MMP-7 expression within EBR tissues from 4 mTLS+ patients. IgG deposition showed a strong positive correlation with MMP-7 expression (Spearman *r* = 0.7744, *P* < 0.0001) (Fig. 5h, i and Supplementary Fig. 5a, b). Using spatial transcriptomic data[9], we further demonstrated that the plasma cell signature[43] and *MMP-7* expression colocalized specifically to fibrotic CK19^+^ bile ducts in BA liver (Fig. 5j). These data are consistent with a model in which TLS-mediated autoantibody production contributes to bile duct injury in BA.

### IL-21R blockade after viral challenge ameliorates biliary atresia in mice

Given the association between IL-21 and autoantibody production observed in our clinical data, we investigated whether targeting IL-21 signaling could ameliorate disease in the rhesus rotavirus (RRV)-infected BA mouse model[44]. Anti-IL-21R antibody was administered beginning 4 days after RRV injection and continued every 48 hours through day 10 (Fig. 6a). Post-infection anti-IL-21R treatment significantly improved multiple disease parameters compared to untreated RRV-induced mice (Fig. 6b). Specifically, treated mice showed improved weight gain (*P* = 0.009, Mann-Whitney U test) (Fig. 6c). Cholangiography demonstrated that IL-21R blockade prevented extrahepatic biliary obstruction (Fig. 6d, e). Histological analysis further revealed patent bile duct lumens in treated animals, whereas untreated RRV-induced mice exhibited marked extrahepatic bile duct wall thickening, luminal hyperplasia, and intraluminal obstruction (Fig. 6f). In addition, post-infection IL-21R blockade reduced portal inflammation (Fig. 6f). Consistent with these histological improvements, serum GGT was significantly reduced in treated animals (*P* = 0.0381, Mann-Whitney U test) (Fig. 6g).

**Fig. 6:**
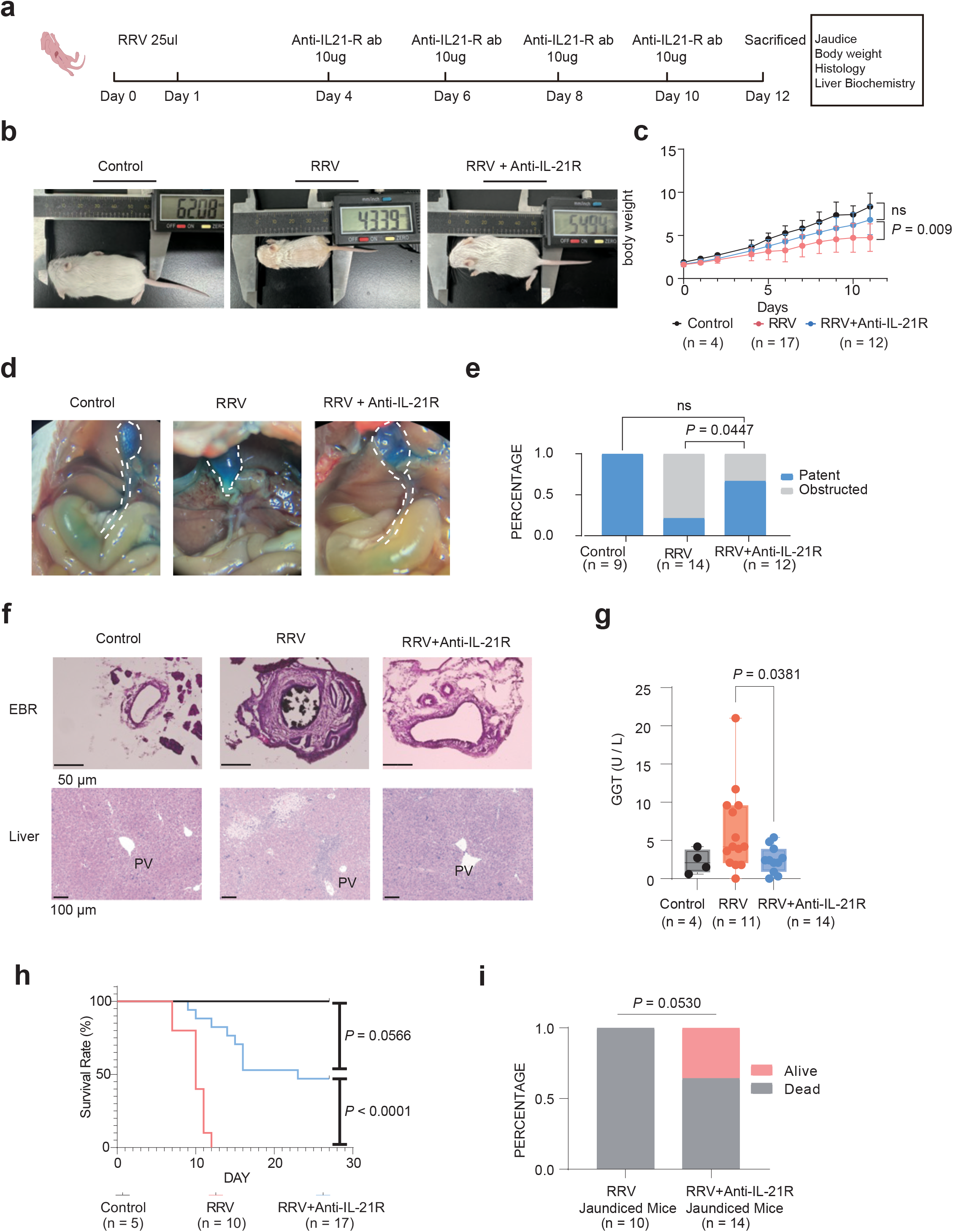
Post-infection anti-IL-21R antibody treatment ameliorates disease phenotype in RRV-induced BA mouse model. (**a**) Experimental design for Anti-IL-21R antibody treatment in rhesus rotavirus (RRV)-infected mice. (**b**) Representative images of mice from different treatment groups. (**c**) Body weight (control, n = 4; RRV-induced, n = 12; RRV-induced mice treated with Anti-IL-21R antibody, n = 17). Statistical analysis was performed using Mann-Whitney U test on day11. (**d**) Representative cholangiography with methylene blue staining. Gallbladder and extrahepatic bile duct are outlined by white dashed lines. (**e**) Bile duct patency rates on day 12 across groups (Fisher’s exact test). (**f**) Representative H&E staining of extrahepatic bile ducts (top) and liver tissues (bottom) from mice in each group. PV, portal vein. (**g**) Serum γ-glutamyl transferase (GGT) levels by treatment group (Student’s t test). **(h)** Kaplan-Meier survival curves comparing 27-day survival among control, RRV, and RRV+Anti-IL-21R groups (log-rank test). **(i)** Mortality rates in jaundiced mice across treatment groups (Fisher’s exact test).

Notably, 4 of 7 jaundiced mice in the treatment group retained patent bile ducts. Comparative histology of the extrahepatic bile ducts and liver in jaundiced mice revealed that treated animals with patent ducts maintained largely preserved ductal architecture, whereas those with obstructed ducts, similar to untreated RRV-induced mice, displayed duct wall thickening. Untreated mice with obstruction also exhibited more extensive inflammatory infiltration in the liver (Supplementary Fig. 6a). The absence of significant differences in serum ALT, AST, and direct bilirubin (DBIL) between treated and untreated groups (Supplementary Fig. 6b) may reflect concurrent cholestatic liver injury in the subset of treated animals that still developed jaundice, suggesting that IL-21R blockade may primarily protect against biliary obstruction rather than hepatocellular injury per se.

Survival analysis further demonstrated that anti-IL-21R antibody treatment not only improved biliary patency but also significantly improved survival in the RRV-induced BA model. By day 27, all control animals survived, whereas the untreated RRV-induced group experienced 100% mortality. In contrast, the anti-IL-21R treatment group achieved a 47% survival rate (8/17), representing a highly significant improvement over the untreated group (*P* < 0.0001, log-rank test) (Fig. 6h). Among mice that developed jaundice, mortality was 100% (10/10) in the untreated group compared with 64% (9/14) in the treatment group (*P* = 0.0530, Fisher’s exact test) (Fig. 6i), indicating that post-infection IL-21R blockade partially improved survival by ameliorating biliary obstruction. Together, these findings establish IL-21/IL-21R signaling as a critical driver of BA pathogenesis and a promising therapeutic target.

## Discussion

BA is a severe neonatal liver disease whose pathogenesis involves progressive liver fibrosis and bile duct obstruction driven, at least in part, by aberrant immune responses. In this study, we provide an integrated characterization of TLSs in BA extrahepatic biliary remnants, identify the enrichment of Tph cells within these structures, and demonstrate that post-infection IL-21R blockade ameliorates disease in a murine BA model. While individual components of the immune landscape in BA have been previously characterized, including lymphocyte infiltration, B cell tolerance defects, and autoantibody production[7,11–13,31], our study adds to this body of work by systematically characterizing TLS maturation states, identifying Tph cells as a disease-associated CD4^+^ T cell subset within these structures, and providing functional evidence for the therapeutic relevance of the IL-21/IL-21R axis.

TLSs are ectopic lymphoid aggregates that form in chronic inflammatory conditions and play crucial roles in sustaining local immune responses. TLSs have been identified in various autoimmune diseases, including rheumatoid arthritis[45], lupus nephritis[46], primary biliary cholangitis[47], and Sjögren’s syndrome[48], where they contribute to local immune activation and tissue destruction. In BA, ectopic lymphoid aggregates in biliary remnants were described by Bove et al.[31], who noted their presence but did not systematically characterize their maturation or clinical significance. Our analysis extends these observations by performing TLS maturation staging in a large cohort (n = 148) and demonstrating that mature TLSs with CD23^+^ germinal centers are associated with elevated disease severity markers. The higher prevalence of TLSs in EBRs compared to liver tissue is consistent with the EBR representing a site of concentrated immune activation, likely reflecting the local antigenic stimulation occurring at the site of initial bile duct injury.

A key finding of our study was the identification and characterization of Tph cells in BA. We designated the CD4^+^PD-1^hi^CXCR5^-^ T cell population as Tph cells rather than Tfh cells based on their absence of CXCR5 expression at both the transcriptomic and protein levels, together with their peripheral localization within TLSs (Fig. 5c). We acknowledge that the nomenclature of these cells remains debated, and that Tph and Tfh cells share substantial functional overlap, including the capacity to promote B cell differentiation via IL-21[38]. Regardless of nomenclature, the functional significance of these Tph cells in providing B cell help within and around TLSs is supported by our ligand-receptor analysis, their spatial co-localization with B cells at TLS peripheries, and the elevated IL-21 levels in the sera of BA patients. These observations parallels findings in systemic lupus erythematosus and IgG4-related disease, in which Tph cells drive pathogenic humoral immunity[49,50]. Although IL-21 can be produced by multiple T cell subsets, including Th17 cells and NK T cells, intracellular IL-21 staining demonstrated that circulating Tph cells in BA produce IL-21 ex vivo. However, the relative contribution of different cellular sources to the systemic IL-21 pool in patients with BA remains to be defined.

Mature TLSs in our cohort were associated with increased rates of autoantibody positivity and localized immunoglobulin deposition on injured bile ducts. This finding aligns with the established role of TLSs in driving local antibody responses in autoimmune diseases[39–41] and extends prior work demonstrating defective B cell tolerance and pathogenic autoantibodies in BA[11,14]. The pathogenic significance of these autoantibodies and their target antigens warrant further investigation using functional assays such as bile duct epithelial organoid models.

Our demonstration that post-infection IL-21R blockade improved biliary patency and long-term survival in the RRV-induced mouse model provides functional evidence that IL-21 signaling contributes to BA disease progression. Anti-IL-21R antibody treatment, initiated on day 4 after viral challenge, significantly reduced jaundice incidence, improved weight gain, and prevented biliary obstruction, ultimately leading to markedly improved long-term survival. Histopathological analysis revealed reduced lymphocyte infiltration in both the liver and extrahepatic bile ducts of treated animals. Notably, improved survival occurred despite similar early elevations in serum bilirubin and liver enzymes between treated and untreated groups, underscoring that maintaining biliary patency, rather than merely mitigating hepatocellular injury, is the critical determinant of survival in this model. These findings position IL-21R blockade as a potential strategy to modify the progression of biliary obstruction in BA.

Several additional limitations of this study merit consideration. Our scRNA-seq analysis, while providing valuable insights into the cellular composition of TLS-containing EBRs, was limited to pooled samples from two BA patients and lacked a non-BA cholangiopathy control group. This limits the generalizability of the findings and our ability to distinguish BA-specific from general cholangiopathy-associated immune responses. The small number of cells in the Tph cluster further necessitates validation in larger, independent datasets. While we used choledochal cyst patients as age-matched controls for bulk RNA-seq and ELISA analyses, these are imperfect controls as they do not fully replicate the cholestatic milieu of BA. Finally, the IgG/IgM deposition analysis was performed in only 4 mTLS+ patients using consecutive tissue sections rather than multiplex immunofluorescence on a single slide, which limits spatial resolution and quantitative rigor.

In summary, our integrated analysis identifies TLSs in BA extrahepatic biliary remnants as organized immune structures harboring disease-associated Tph cells that are associated with elevated IL-21, autoantibody production, and bile duct injury markers. Post-infection IL-21R blockade ameliorated biliary obstruction and improved survival in a murine BA model. These findings support a model in which TLS-associated Tph cells contribute to BA pathogenesis through IL-21-mediated B cell activation and suggest that the IL-21/IL-21R axis warrants further investigation as a potential therapeutic target.

## Data Availability

All other data, analytic methods, and study materials will be made available to other researchers upon reasonable request.

## Acknowledgements

This study was supported by the National Natural Science Foundation of China (No. 82270541, and 32400835), the National Clinical Key Specialty Construction Project (No. 10000015Z155080000004), the Shanghai Municipal Key Clinical Specialty (No. shslczdzk05703), the National Children’s Medical Center Projects (No. EKYX202401, EKSJD202409 and EKQM202402), and the China Postdoctoral Science Foundation (No. 2023M730687), Shanghai Municipal Health Commission (no. 2025ZZ2018).

## Abbreviations

ANA: antinuclear antibody
BA: biliary atresia
CC: choledochal cyst
EBR: extrahepatic biliary remnant
ELISA: enzyme-linked immunosorbent assay
GGT: γ-glutamyl transferase
H&E: hematoxylin and eosin staining
IHC: immunohistochemistry
IL-21: interleukin-21
mIHC: multiplex immunohistochemistry
mTLS: mature tertiary lymphoid structure
PBMC: peripheral blood mononuclear cell
RRV: rhesus rotavirus
scRNA-seq: single-cell RNA sequencing
ST: spatial transcriptomics
Tfh: follicular helper T cell
TLS: tertiary lymphoid structure
Tph: T peripheral helper cell
UMAP: Uniform Manifold Approximation and Projection.

## Author contributions

Conceptualization: GC and YL

Methodology: MTL, WDM, and YL

Investigation: MTL, WDM, YKC, SJW, MNQ, DC, JZ, JXD, YFY, JJY, TL, QS, and XFG

Visualization: MTL and WDM

Funding acquisition: WDM, SZ, GC, and YL

Project administration: YL and GC

Supervision: YL, GC, XMZ, SZ, RD, WQQ, and SYS

Writing - original draft: MTL and WDM

Writing - review & editing: MTL, WDM, XMZ, GC, and YL

