## Supplementary Figure 1 for "IL-21-producing peripheral helper T cells associate with autoimmune bile duct injury in biliary atresia"

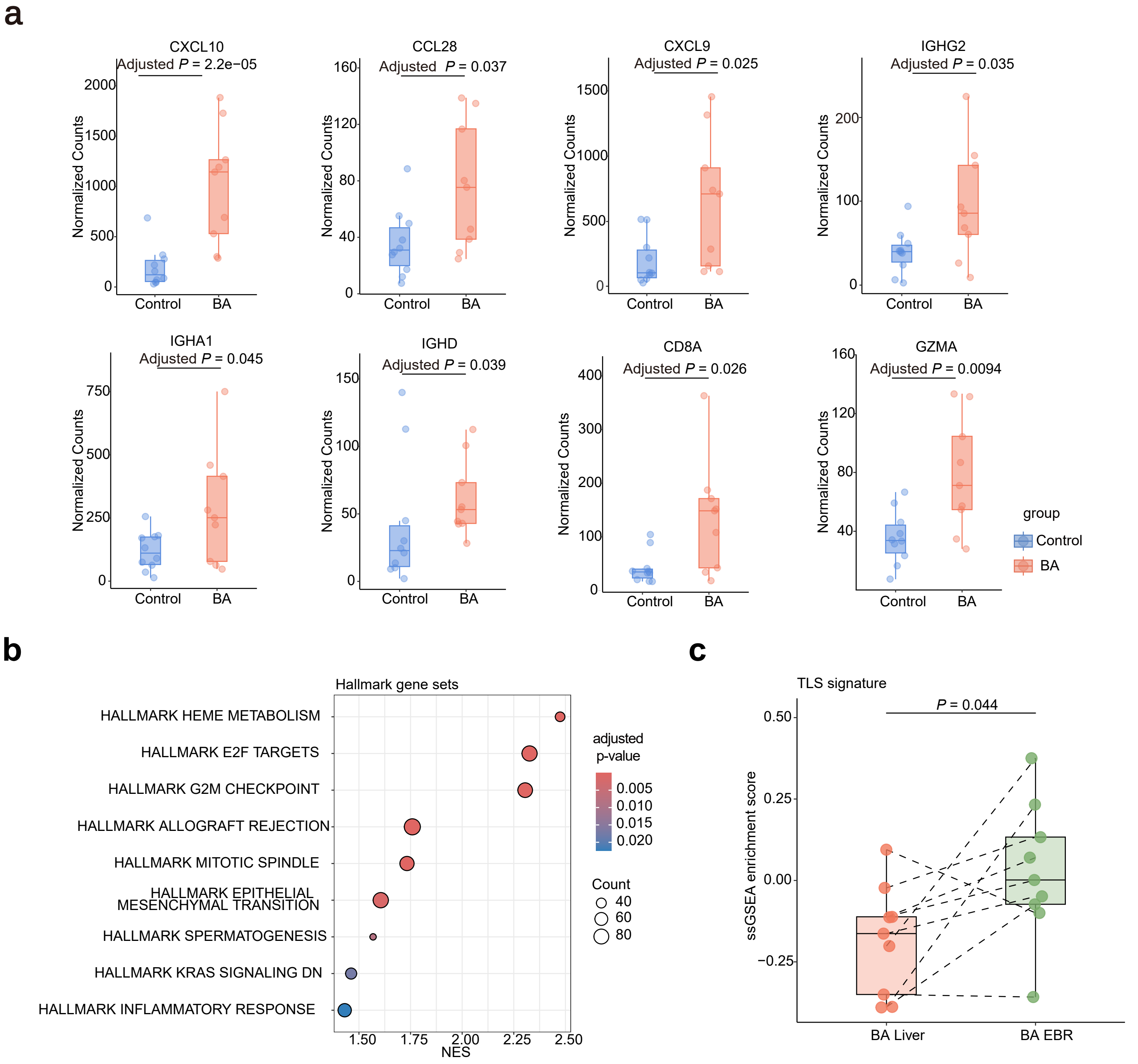


**Supplementary Figure 1: Activation of immune response pathways in BA liver and EBR.** (**a**) Box plots depicting the mRNA expression levels of the indicated genes in BA liver (n = 9) versus control liver tissues (n = 10). Adjusted P-values are shown at the top. (**b**) Bubble chart showing significant GSEA results for hallmark gene sets comparing BA liver with Control liver. Pathways highlighted in bold represent those associated with inflammatory responses. (**c**) Dot plot with connecting dashed lines showing TLS signature scores in BA EBRs versus paired liver tissues (n = 9) (paired Student's t test), estimated using the ssGSEA algorithm.
