## Supplementary Figure 2 for "IL-21-producing peripheral helper T cells associate with autoimmune bile duct injury in biliary atresia"

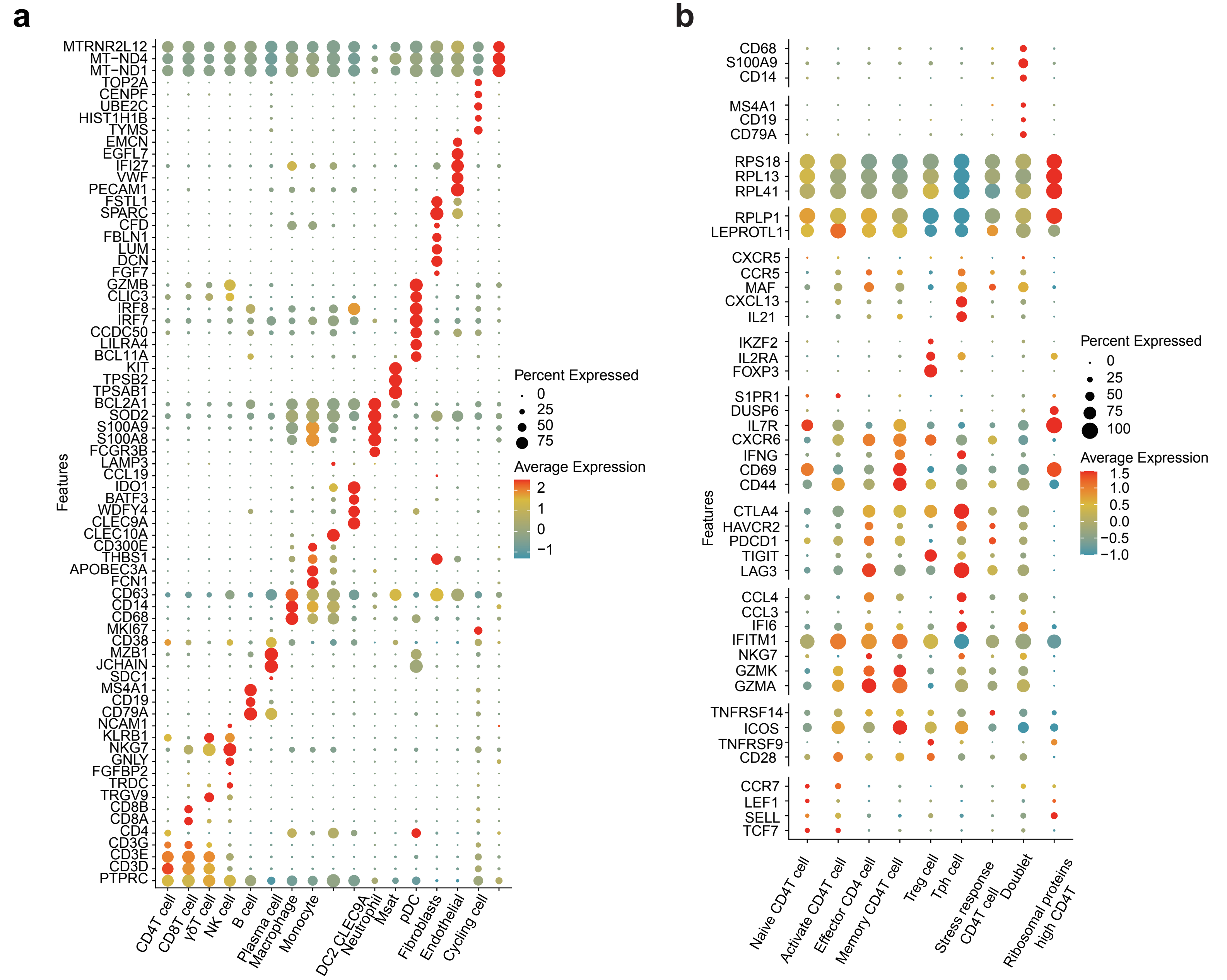


**Supplementary Figure 2: Expression of marker genes in specific cell subsets.** (**a-b**) Dotplots depicting canonical marker gene expression in 17 clusters (**a**) and 9 $CD4^{+}$ T subtypes (**b**) obtained from single-cell RNA sequencing.
