## Supplementary Figure 3 for "IL-21-producing peripheral helper T cells associate with autoimmune bile duct injury in biliary atresia"

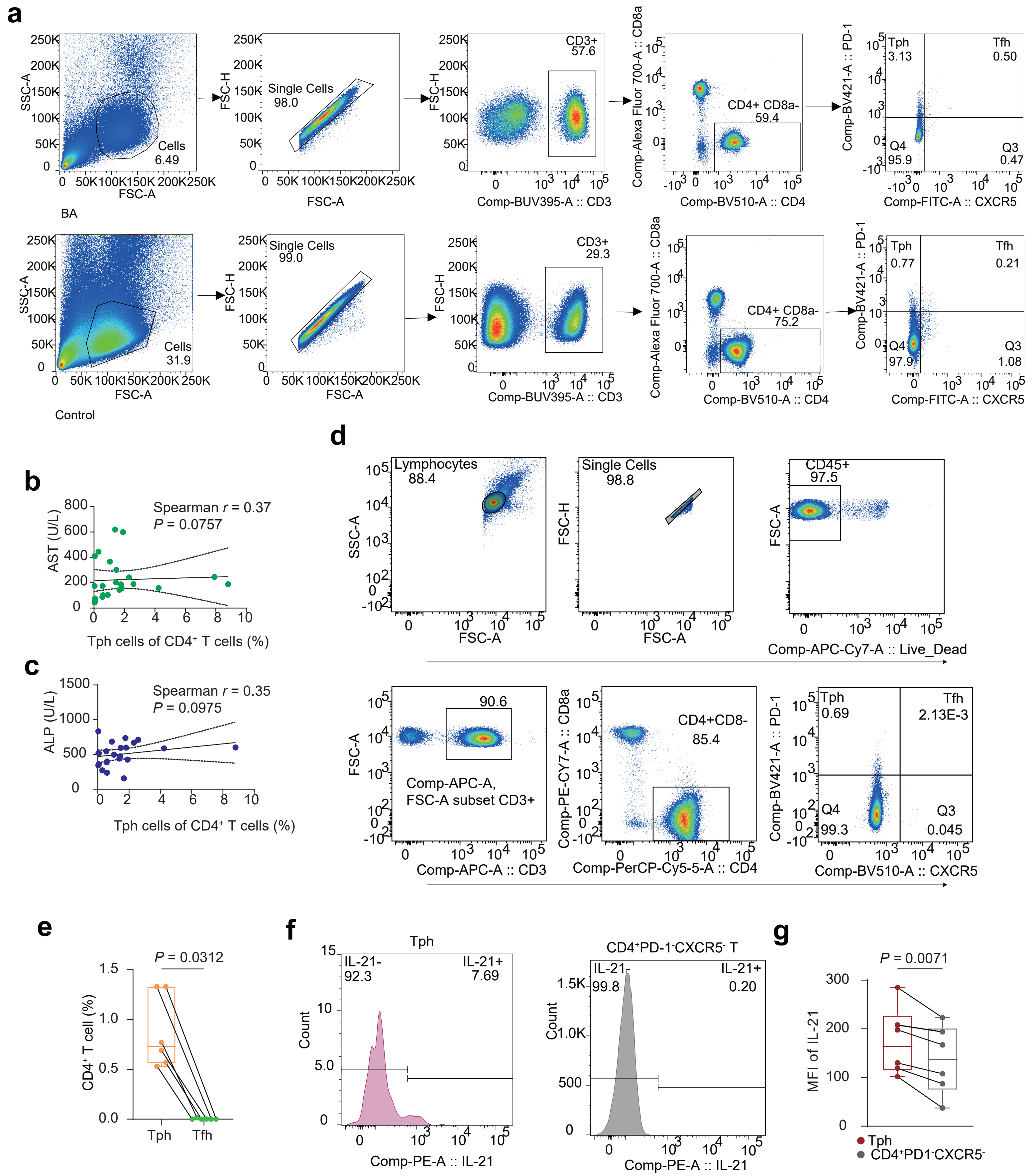

**Supplementary Figure 3: Comparative analysis of peripheral blood Tph and Tfh cells in biliary atresia.** (**a**) Gating strategy for flow cytometric identification of CD3⁺CD4⁺CD8a⁻PD-1ʰⁱCXCR5⁻ Tph cells in PBMCs from patients with BA (upper panel) and age-matched non-BA infantile cholestasis controls (lower panel). (**b**) Correlation between the frequency of peripheral Tph cells and serum AST levels in BA patients (n=18) and age-matched non-BA cholestasis controls (n=6), analyzed by Spearman correlation. (**c**) Correlation between the frequency of peripheral Tph cells and serum ALP levels in BA patients (n=17) and age-matched non-BA cholestasis controls (n=6), analyzed by Spearman correlation. (**d**) Gating strategy for flow cytometric analysis of CD45⁺CD3⁺CD4⁺CD8α⁻PD-1ʰⁱCXCR5⁻ Tph cells in BA PBMCs following in vitro stimulation with BD Leukocyte Activation Cocktail containing GolgiPlug™. (**e**) Frequencies of Tfh (CD45⁺CD3⁺CD4⁺CD8α⁻PD-1ʰⁱCXCR5⁺ ) and Tph cells in BA PBMCs (n=6) (Wilcoxon signed-rank test). (**f**) Representative flow cytometry histograms showing IL-21 expression in Tph cells and CD45⁺CD3⁺CD4⁺CD8a⁻PD-1⁻CXCR5⁻ T cells from BA patients (n=6). (**g**) IL-21 mean fluorescence intensity (MFI) in Tph cells vs. CD45⁺CD3⁺CD4⁺CD8a⁻PD-1⁻CXCR5⁻ T cells (Wilcoxon signed-rank test).
