## Supplementary Figure 4 for "IL-21-producing peripheral helper T cells associate with autoimmune bile duct injury in biliary atresia"

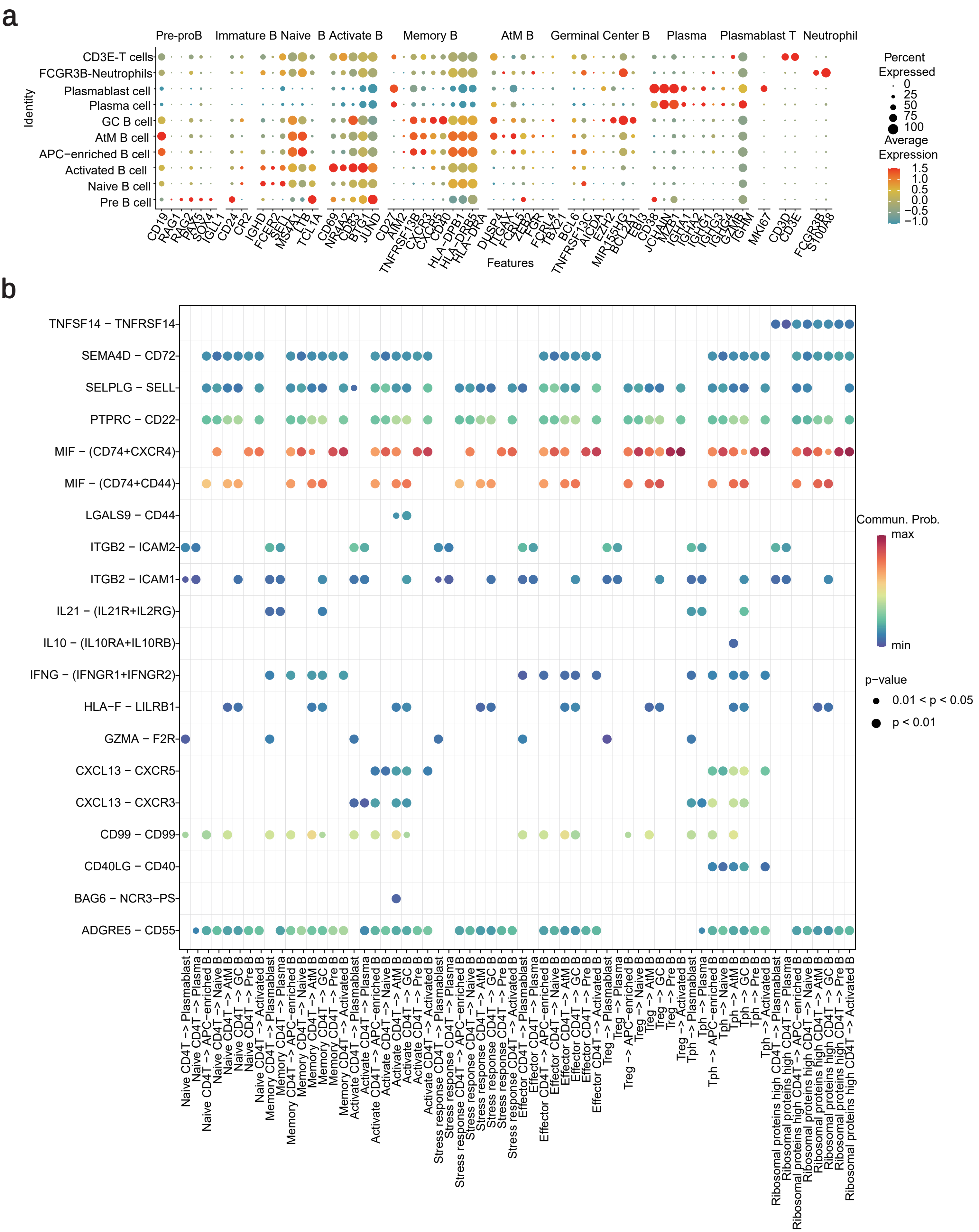


**Supplementary Figure 4: Marker gene expression in B cell subsets and ligand-receptor interaction analysis in scRNA-seq.** (**a**) Dot plot depicting canonical B cell subset and plasma cell marker genes expression. (**b**) Bubble chart showing all significant interactions (ligand-receptor pairs) from CD4$^{+}$T cells to B cells and plasma cells.
