## Supplementary Figure 5 for "IL-21-producing peripheral helper T cells associate with autoimmune bile duct injury in biliary atresia"

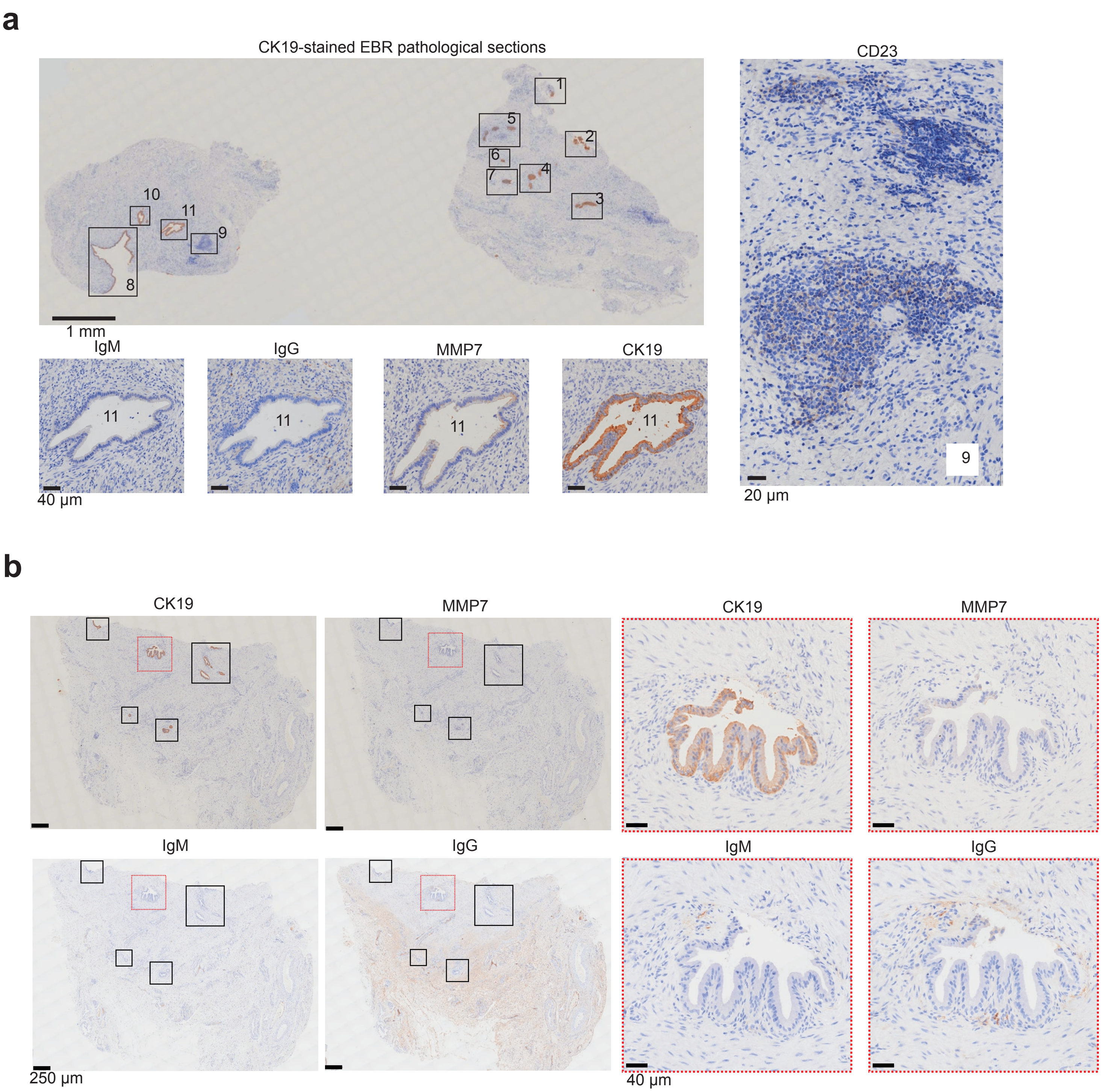


**Supplementary Figure 5: Autoantibodies and In-Situ Deposition in Biliary Atresia.** (**a-b**) Representative immunohistochemical images of consecutive tissue sections showing CK19, MMP-7, anti-human IgM, and anti-human IgG expression in EBR tissue from a BA patient with mTLS+ (**a**) and mTLS- (**b**).
