## Supplementary Figure 6 for "IL-21-producing peripheral helper T cells associate with autoimmune bile duct injury in biliary atresia"

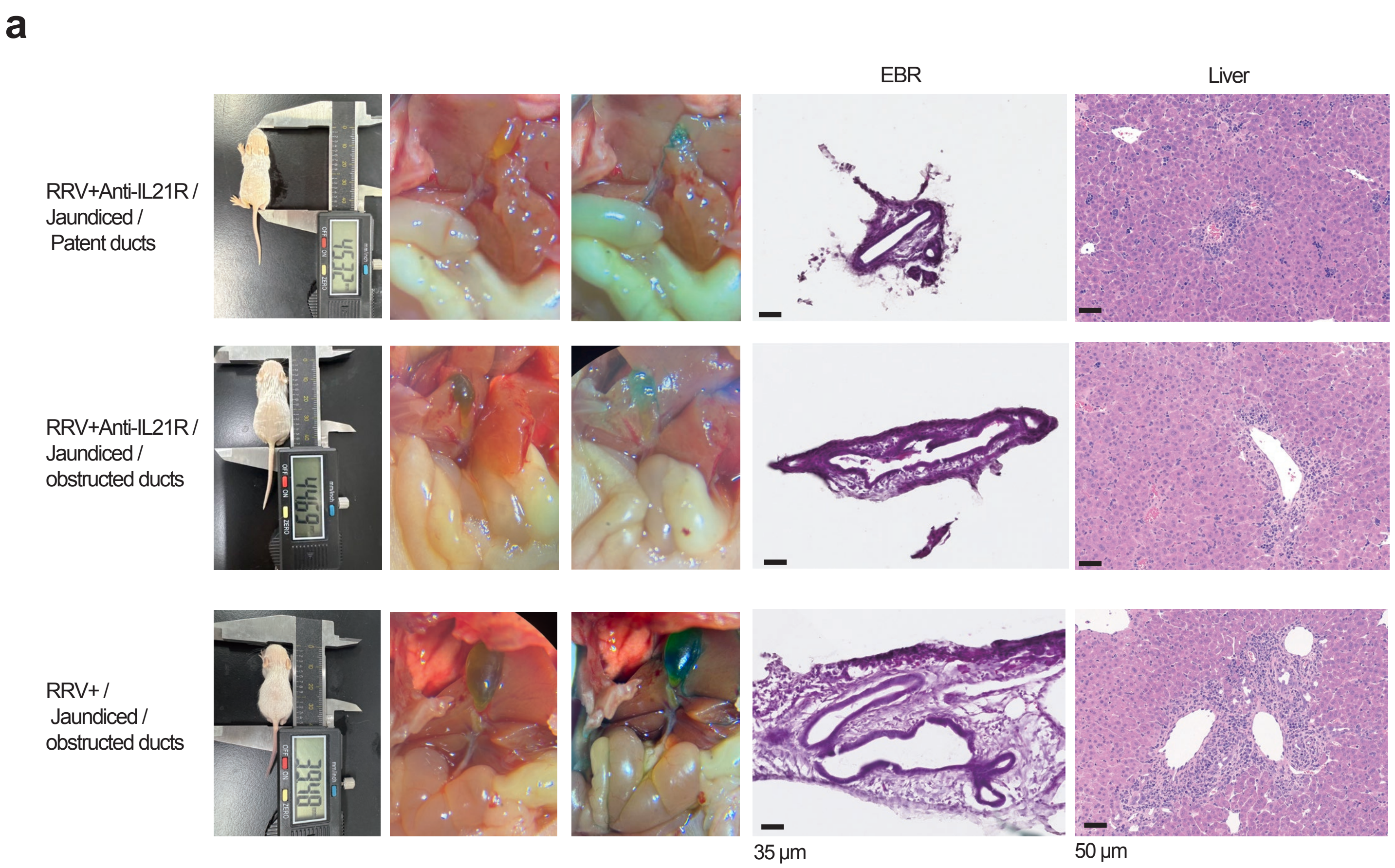


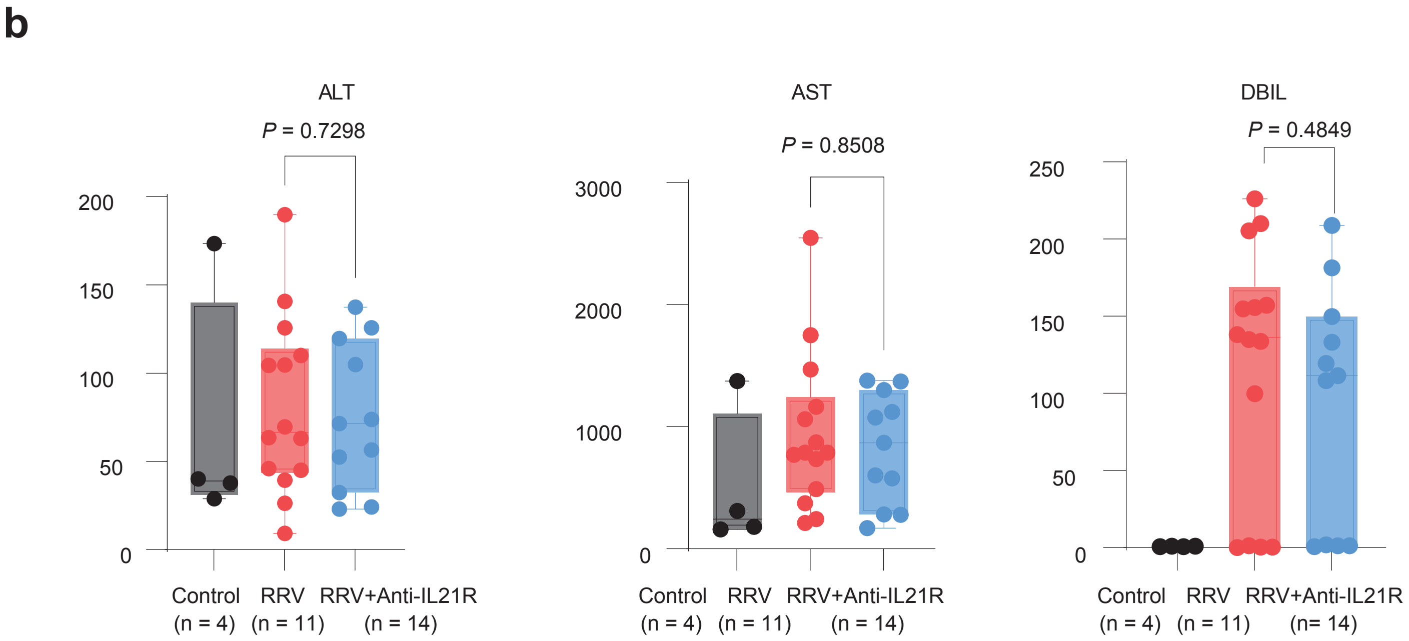


**Supplementary Figure 6: Biliary phenotype and associated liver function biomarkers in the BA model.** (**a**) Representative images illustrating the gross appearance, along with methylene blue and H&E staining of the extrahepatic bile ducts and liver tissues, are presented for the different experimental groups. (**b**) Serum levels of alanine aminotransferase (ALT), aspartate aminotransferase (AST) and direct bilirubin (DBIL) in each treatment group (Student’s t test).
