## Supplementary Table1 for "IL-21-producing peripheral helper T cells associate with autoimmune bile duct injury in biliary atresia"

**table S1: Canonical marker genes used to annotate 17 clusters from CD45^+^ cells**

| **Cell clusters** | **Marker genes** | | | | | | | | **References** |
| --- | --- | --- | --- | --- | --- | --- | --- | --- | --- |
| CD45 | *PTPRC* |  |  |  |  |  |  |  | 1. Maynard A, et al. ***Cell*** (2020) 182(5):1232-1251.e22 2. Zhang R, et al. ***J Hepatol*** (2022) 77(5):1299-1310 3. Wang J, et al. ***Cell*** (2020) 183(7):1867-1883.e26. 4. Ye C, et al. ***Clin Transl Med*** (2022) 12(11):e1070 |
| T cell | *CD3D* | *CD3E* | *CD3G* | *CD4* | *CD8A* | *CD8B* | *TRGV9* | *TRDC* |  |
| NK cell | *FGFBP2* | *FCG3RA* | *GNLY* | *NKG7* | *KLRB1* | *NCAM1* |  |  |  |
| B cell | *CD79A* | *CD19* | *MS4A1* |  |  |  |  |  |  |
| Plasma | *SDC1* | *JCHAIN* | *MZB1* |  |  |  |  |  |  |
| Plasmablast | *CD38* | *MKI67* |  |  |  |  |  |  |  |
| Macrophage | *CD68* | *CD14* | *CD63* |  |  |  |  |  |  |
| Monocyte | *FCN1* | *APOBEC3A* | *THBS1* | *CD300E* |  |  |  |  |  |
| DC | *CLEC10A* | *CLEC9A* | *WDFY4* | *BATF3* | *IDO1* | *CCL19* | *LAMP3* |  |  |
| Neutrophil | *FCGR3B* | *S100A8* | *S100A9* | *SOD2* | *BCL2A1* |  |  |  |  |
| Mast | *TPSAB1* | *TPSB2* | *KIT* |  |  |  |  |  |  |
| pDC | *BCL11A* | *LILRA4* | *CCDC50* | *IRF7* | *IRF8* | *CLIC3* | *GZMB* |  |  |
| Fibroblasts | *FGF7* | *DCN* | *LUM* | *CD10* | *FBLN1* | *CFD* | *SPARC* | *FSTL1* |  |
| Endothelial | *PECAM1* | *VWF* | *CD31* | *IFI27* | *EGFL7* | *EMCN* |  |  |  |
| Cycling_cell | *TYMS* | *HIST1H1B* | *UBE2C* | *CENPF* | *TOP2A* |  |  |  |  |
| Mitochondrial_gene | *MT-ND1* | *MT-ND4* | *MTRNR2L12* |  |  |  |  |  |  |
