## Supplementary Table2 for "IL-21-producing peripheral helper T cells associate with autoimmune bile duct injury in biliary atresia"

| **Cell clusters** | **Marker genes** | | | | | | | | | **References** |
| --- | --- | --- | --- | --- | --- | --- | --- | --- | --- | --- |
| Naïve CD4^+^T | *TCF7*^+^ | *SELL*^+^ | *LEF1*^+^ | *CCR7*^+^ |  |  |  |  |  | 1. Xiong, Jie et al. ***BMC Biology*** (2023) 21(1):46 2. Bocharnikov AV, et al. ***JCI Insight*** (2019) 4(20):e130062. 3. Ji Z, et al. ***Int J Mol Sci***. (2023) 24(18):13735 4. Wang J, et al. ***Cell*** (2020) 183(7):1867-1883.e26 |
| Activate CD4^+^T | *CD28^+^* | *TNFRSF9^+^* | *ICOS^+^* | *TNFRSF14^+^* | *CD62^+^* |  |  |  |  |  |
| Effector CD4^+^T cells | *GZMA*^+^ | *PRF1*^+^ | *GZMB*^+^ | *GZMK*^+^ | *IFNG*^+^ | *NKG7*^+^ | *IFI6*^+^ | *IFITM1*^+^ | *IFI27L2*^+^ |  |
| Memory CD4^+^T | CD44^+^ | CD69^+^ | IFNG^+^ | CXCR6^+^ | IL7R^+^ | DUSP6^+^ | S1PR1^+^ |  |  |  |
| Treg cells | *FOXP3^+^* | *IL2RA^+^* | *IKZF2^+^* |  |  |  |  |  |  |  |
| Tph cells | *PDCD1*^+^ | *TIGIT*^+^ | *ICOS*^+^ | *CXCR5*^-^ | *CXCL13*^+^ | *IL21*^+^ | *IFN-γ*^+^ | *MAF*^+^ |  |  |
| Stress response CD4T^+^ cells | *RPLP1^+^* | *LEPROTL1^+^* |  |  |  |  |  |  |  |  |
| Doublet | *CD68^+^* | *S100A9^+^* | *CD14^+^* | *MS4A1^+^* | *CD19^+^* | *CD79A^+^* |  |  |  |  |
| Ribosomal proteins-high CD4^+^ T cells | *RPL41*^+^ | *RPL13*^+^ | *RPS18*^+^ |  |  |  |  |  |  |  |

**table S2: Canonical marker genes used to annotate 9 clusters from CD4^+^ T cells**
