## Supplementary Table3 for "IL-21-producing peripheral helper T cells associate with autoimmune bile duct injury in biliary atresia"

| **B cell subsets** | **Marker genes** | | | | | | | | | **References** |
| --- | --- | --- | --- | --- | --- | --- | --- | --- | --- | --- |
| pre-B | *CD19^+^* | *CD34^-^* | *RAG1^+^* | *RAG2^+^* | *PAX5^+^* | *SOX4^+^* | *IGLL1^+^* |  |  | 1. Wang J, et al. ***Cell*** (2020) 183(7):1867-1883.e26 2. Ma J, et al. ***Science*** (2024) 384(6695):eadj4857 |
| naïve B | *IGHD^+^* | *FCER2^+^* | *SELL^+^* |  |  |  |  |  |  |  |
| activated B | *CD69^+^* | *NR4A2^+^* | *CD83^+^* | *BTG1^+^* | *JUND^+^* |  |  |  |  |  |
| APC-enriched B | *HLA-DPB1^+^* | *HLA-DRB5^+^* | *HLA-DRA^+^* |  |  |  |  |  |  |  |
| atypical memory B | *CR2^-^* | *CD27^-^* | *IgD^-^* | *DUSP4^+^* | *ITGAX^+^* | *FCRL5^+^* | *ZEB2^+^* | *FGR^+^* | *FCRL4^+^* |  |
| germinal center B | *BCL6^+^* | *AICDA^+^* | *EZH2^+^* | *MIR155HG^+^* | *BCL2A1^+^* | *EBI3^+^* |  |  |  |  |
| plasma cell | *CD38^+^* | *JCHAIN^+^* | *MZB1^+^* |  |  |  |  |  |  |  |
| plasmablasts | *CD38^+^* | *JCHAIN^+^* | *MZB1^+^* | *MKI67^+^* |  |  |  |  |  |  |
| FCGR3B-Neutrophils | *FCGR3B^+^* | *S100A8^+^* |  |  |  |  |  |  |  |  |
| CD3E-T cells | *CD3D^+^* | *CD3E^+^* |  |  |  |  |  |  |  |  |

**table S3: Canonical marker genes used to annotate 10 B cell subsets**
