## Supplementary Table4 for "IL-21-producing peripheral helper T cells associate with autoimmune bile duct injury in biliary atresia"

| Samples used in RNA-seq analysis | | | | |
| --- | --- | --- | --- | --- |
|  | Characteristics | BA (Type III)  (n = 9) | Choledochal cysts  (n = 10) | *P* value |
|  | Age, median (IQR), day | 68(53.5-89.5) | 173.5(52-355.5) | 0.1128 |
|  | Sex, No. (%) |  |  | 0.6285 |
|  | Male | 3(33.3%) | 2(20%) |  |
|  | Female | 6(66.7%) | 8(80%) |  |
|  | TBIL, median (IQR) | 139.9(109.1-163.1) | 10.4(6.3-98.95) | **0.0315** |
|  | DBIL, mean (SD) | 101.7(26.01) | 5.656(4.042) | **<0.0001** |
|  | ALT, median (IQR) | 82.77(59.34-115.7) | 26.78(15.92-60.26) | **0.0188** |
|  | AST, median (IQR) | 153.2(100.5-214.9) | 41.36(35.54-84.34) | **0.004** |
|  | ALP, median (IQR) | 519.5(372.5-622.7) | 289.1(173.3-356.4) | **0.0188** |
|  | GGT, median (IQR) | 290.9(210.3-458.1) | 21.85(13.76-674.6) | 0.2224 |
|  | ALB, mean (SD) | 37.99(2.743) | 40.92(5.304) | 0.1606 |
|  | PALB, mean (SD) | 148.9(53.8) | 163.5(39.76) | 0.5204 |
|  | TBA, mean (SD) | 86.4(20.34) | 12.7(10.42) | **<0.0001** |

**table S4: Sample characteristics**

| BA samples (type III) used for histological staining | | |
| --- | --- | --- |
|  | Characteristics | Values  (n = 148) |
|  | Age, median (IQR), day | 57(41-69) |
|  | Sex, No. (%) |  |
|  | Male | 69(46.6%) |
|  | Female | 79(53.4%) |
|  | TBIL, median (IQR) | 149.3(124.9-189.9) |
|  | DBIL, median (IQR) | 117(93.75-151.3 |
|  | ALT, median (IQR) | 125.7(71.65-191.2) |
|  | AST, median (IQR) | 180.9(122.6-267.9) |
|  | ALP, median (IQR) | 490(371-647.5) |
|  | GGT, median (IQR) | 373.6(209.7-660.4) |
|  | ALB, median (IQR) | 38.23(35.6-40.38) |
|  | PALB, median (IQR) | 110(85-142) |
|  | TBA, median (IQR) | 90.7(73.05-108.9) |

| Samples used in flow cytometry analysis | | | | |
| --- | --- | --- | --- | --- |
|  | Characteristics | BA (Type III)  (n = 18) | Choledochal cysts  (n = 6) | *P* value |
|  | Age, median (IQR), day | 59(52-71) | 50(38-69) | 0.449 |
|  | Sex, No. (%) |  |  | >0.9999 |
|  | Male | 5(27.8%) | 1(16.67%) |  |
|  | Female | 13(72.2%) | 5(83.33%) |  |
|  | TBIL, median (IQR) | 163.9(124.1-201.8） | 105(43.9-150.6） | **0.0143** |
|  | DBIL, median (IQR) | 125.3(98.75-141.9） | 35.5(9.925-89.73） | **0.0004** |
|  | ALT, median (IQR) | 124.1(90.65-135) | 28.74(23.28-162.7) | 0.0524 |
|  | AST, median (IQR) | 188.4(152.9-317.7) | 88.25(48.71-241.9) | **0.0472** |
|  | ALP, median (IQR) | 587(456.5-676.9) | 372.9(324.9-427.5) | **0.0126** |
|  | GGT, median (IQR) | 560.2(385.4-1109) | 315(214.9-600.4) | 0.2011 |
|  | ALB, median (IQR) | 39.31(36.38-42.96) | 37.11(33.46-38.08） | 0.1009 |
|  | PALB, median (IQR) | 99(74.35-118) | 111.3(73.85-142.5) | 0.7506 |
|  | TBA, median (IQR) | 97(79.03-121.5) | 46.8(8.5-75.75) | **0.0036** |

| Samples used in ELISA | | | | |
| --- | --- | --- | --- | --- |
|  | Characteristics | BA (Type III)  (n = 74) | non-BA infantile cholestasis controls  (n = 19) | *P* value |
|  | Age, mean (SD), day | 69.74(24.13) | 78.84(43.01) | 0.2235 |
|  | Sex, No. (%) |  |  | 0.3525 |
|  | Male | 34(46%) | 11(57.9%) |  |
|  | Female | 40(54%) | 8(42.1%) |  |
|  | TBIL, mean (SD) | 161.7(45.5) | 109.3(71.85) | **0.0002** |
|  | DBIL, median (IQR) | 111.7(85.58-135.2) | 75(7.6-120.1) | **0.0037** |
|  | ALT, median (IQR) | 100.1(61.26-212) | 139.6(59.59-206.3) | 0.7089 |
|  | AST, median (IQR) | 174.7(99-282.3) | 144(107.1-270.5) | 0.4965 |
|  | ALP, median (IQR) | 514.3(409.2-671.2) | 442.9(319.4-658.2) | 0.3083 |
|  | GGT, median (IQR) | 438.8(280.9-663.9) | 168.6(63.67-473.5) | **0.0022** |
|  | ALB, mean (SD) | 38.07(3.478) | 39.58(4.648) | 0.1338 |
|  | PALB, median (IQR) | 107(82.75-135) | 151(111.5-193.1) | **0.0031** |
|  | TBA, median (IQR) | 92.95(77.43-115) | 96.25(17.63-210.7) | 0.9855 |

| Samples used in ANA analysis | | |  | |
| --- | --- | --- | --- | --- |
|  | Characteristics | BA (Type III)  (n = 54) | non-BA infantile cholestasis controls  (n = 10) | *P* value |
|  | Age, median (IQR),day | 53(38.5-64.5） | 120.5(37-435.8) | 0.1074 |
|  | Sex, No. (%) |  |  | 0.3268 |
|  | Male | 26(48.15%) | 3(30%) |  |
|  | Female | 28(51.85%) | 7(70%) |  |
|  | TBIL, median (IQR) | 156.7(127.3-187.6) | 41.35(10.7-120.6) | **0.0002** |
|  | DBIL, median (IQR) | 117.1(87.8-140.1) | 9.85(4.95-43.83) | **<0.0001** |
|  | ALT, median (IQR) | 114.2(67.34-169) | 26.93(22.98-167.5) | **0.0286** |
|  | AST, median (IQR) | 170.9(118.6-260.4) | 44.69(39.38-184.6) | **0.0038** |
|  | ALP, median (IQR) | 497(411.3-604.1) | 351.1(311.2-5499) | 0.3573 |
|  | GGT, median (IQR) | 420.8(236.5-715) | 385(51.35-888.7) | 0.555 |
|  | ALB, mean (SD) | 38.74(2.719) | 40.50(4.72) | 0.1039 |
|  | PALB, median (IQR) | 105(84.8-138.3) | 142.7(121.6-184.3) | **0.0071** |
|  | TBA, median (IQR) | 89.55(71.53-102.8) | 12.5(6.5-68.9) | **0.0009** |
